# Alternate conformational trajectories in protein synthesis

**DOI:** 10.1101/2024.04.03.588007

**Authors:** Jose L. Alejo, Dylan Girodat, Michael J. Hammerling, Jessica A. Willi, Michael C. Jewett, Aaron E. Engelhart, Katarzyna P. Adamala

**Affiliations:** Department of Genetics, Cell Biology and Development, University of Minnesota, Minneapolis, MN, USA; Department of Chemistry and Biochemistry, University of Arkansas, Fayetteville, AR, USA; Department of Chemical and Biological Engineering, Northwestern University, Evanston, IL, USA; Department of Bioengineering, Stanford University, Stanford, CA, USA

## Abstract

Translocation in protein synthesis entails the efficient and accurate movement of the mRNA-[tRNA]_2_ substrate through the ribosome after peptide bond formation. An essential conformational change during this process is the swiveling of the small subunit head domain about two rRNA ‘hinge’ elements. Using directed evolution and molecular dynamics simulations, we derive alternate hinge elements capable of translocation *in vitro* and *in vivo* and describe their effects on the conformational trajectory of the EF-G-bound, translocating ribosome. In these alternate conformational pathways, we observe a diversity of swivel kinetics, hinge motions, three-dimensional head domain trajectories and tRNA dynamics. By finding alternate conformational pathways of translocation, we identify motions and intermediates that are essential or malleable in this process. These findings highlight the plasticity of protein synthesis and provide a more thorough understanding of the available sequence and conformational landscape of a central biological process.

**Author Summary:** Translocation, the motion of the ribosome across its mRNA substrate, is an essential stage of protein synthesis. A key conformational change in this process is the rotation of the ribosome head domain about two rRNA hinges in the direction of translocation, repositioning the mRNA and tRNAs in their final states. Employing directed evolution, we obtain variant hinges capable of performing translocation in vitro and in vivo. Through molecular dynamics simulations, the different variant ribosome translocation conformational trajectories are described. This description reveals different possible conformational pathways to translocation, with varying dynamics, motions and intermediates. The understanding of this conformational malleability can increase our knowledge of protein synthesis function, disruption, evolution, and engineering.

## Introduction

Conformational changes in macromolecules are changes in structure that are inextricably linked to signaling, cellular locomotion, and catalysis. The exploration of the inherent flexibility and adaptability accessible to macromolecules during conformational rearrangements is important to understanding their functionality, evolution and engineering. In this study, directed evolution and molecular dynamics were used to address these questions for the process of ribosome translocation.

Translocation, the motion of the mRNA and tRNAs through the ribosome, is an essential macromolecular conformational change central to all of life (1–5). The conserved GTPase EF-G (**Fig. 1a**) accelerates this process, directionally moving the mRNA-[tRNA]_2_ substrates through the aminoacyl (A), peptidyl (P) and exit (E) binding sites formed by the large (50S in prokaryotes, LSU) and small (30S in prokaryotes, SSU) ribosomal subunits. EF-G(GTP) engages the pre-translocation (PRE) complex containing deacylated tRNA in the P site and peptidyl tRNA in the A site, facilitating the rapid and accurate movement of the mRNA-[tRNA]_2_ substrate by one codon (**Fig. 1b**). Structural (6–11), ensemble (12–14) and single-molecule (7, 15–17) imaging studies have established a global model of ribosome translocation. This framework includes three identified intermediate steps (INT1-3) with distinct conformations (7, 15) (**Fig. 1c**). EF-G(GTP) initially binds the rotated conformation of the PRE complex, where the SSU has rotated ∼8°-10° with respect to the LSU and the deacylated P-site tRNA is in a hybrid (P/E) conformation (15, 17). Engagement of the A-site leads EF-G to catalyze GTP hydrolysis, causing ‘unlocking’ (13, 18) and generating various translocation intermediates on the path to the post-translocation (POST) complex, involving the motion of the LSU, tRNAs and SSU head, platform, and body domains (6, 7, 9).

**Figure 1.**
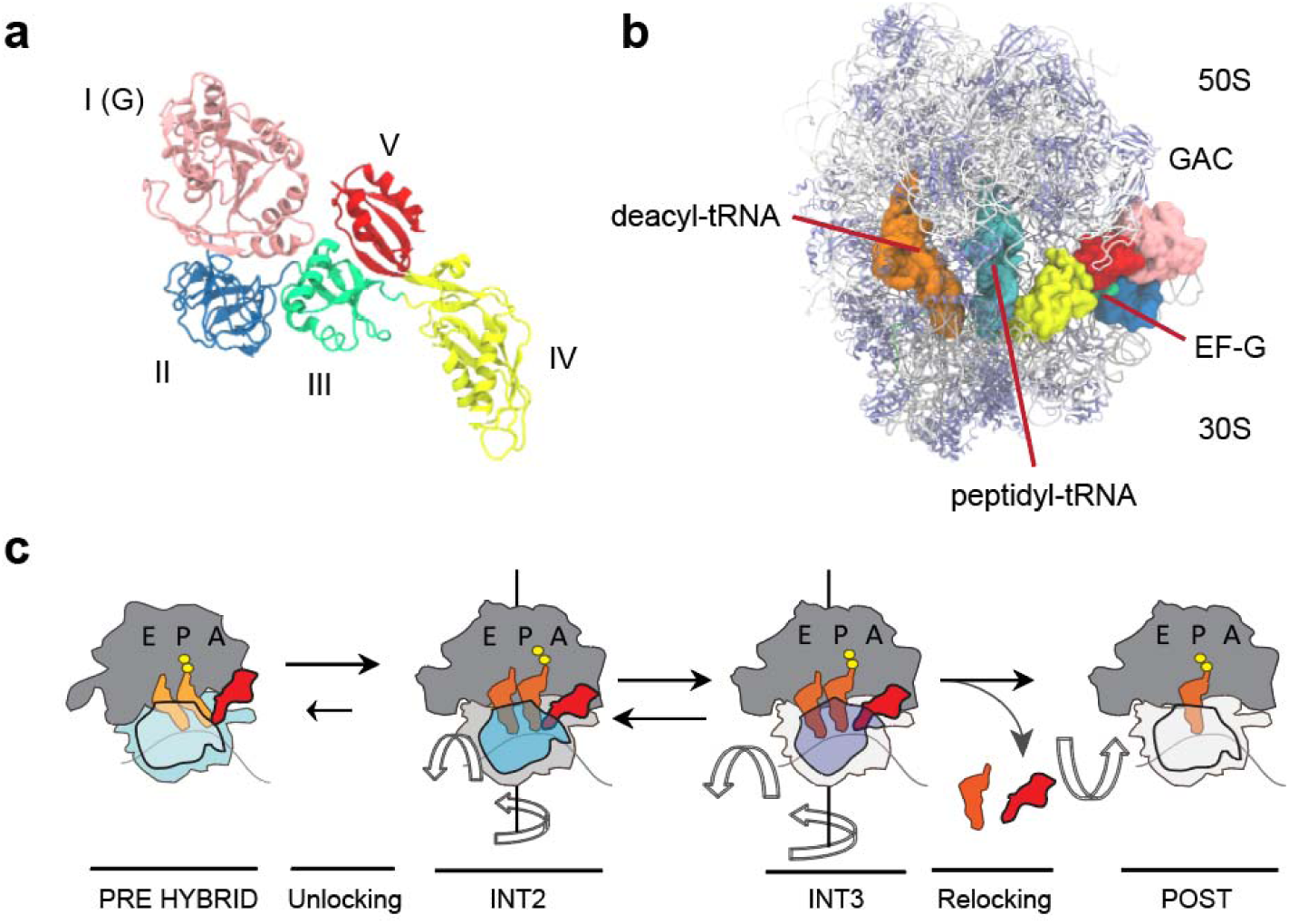
Bacterial ribosome and EF-G structures and translocation scheme. (**a**) EF-G domains I–V are indicated. This structural model is based on Protein Data Bank (PDB) ID code 4V7B (Ref. 9). (**b**) EF-G–bound, FA-stalled PRE complex showing compacted positions of deacylated and peptidyl-tRNAs (orange). The rRNA is shown in gray, the ribosomal proteins are shown in blue. The GTPase activating center (GAC) and EF-G are indicated. The structural model is based on PDB ID code 4W29 (Ref. 8). (**c**) Framework of ribosome translocation in which EF-G(GTP) binds preferentially to PRE complexes in which the small subunit (blue) has rotated with respect to the large subunit (dark gray). EF-G–catalyzed GTP hydrolysis induces “unlocking”, a process that allows the mRNA and tRNA substrates to move relative to the small subunit. Unlocking promotes the (mostly irreversible) formation of the INT2 complex, characterized by tRNA compaction, ∼15°–17° head swivel (blue), and partial small subunit back-rotation (light gray). Further (∼23°–26°) head swivel (purple), together with complete small subunit back-rotation (white) lead to the (reversible) formation of the INT3 state. Recognition of peptidyl tRNA in the small subunit P site stabilizes INT3 and prompts reverse swivel of the head domain, returning it to its original position (white), from which EF-G(GDP) dissociates.

Initially, the EF-G(GDP)-bound ribosome adopts INT1, a spectinomycin-stabilized (19), rapidly transited (15, 20) state consisting of complete and partial translocation of deacyl- and peptidyl-tRNA with respect to the LSU respectively; complete rotation of the SSU and partial swivel of the head domain relative to the SSU body in the direction of translocation (7). This state is followed by a more stable intermediate, INT2, exhibiting partially reversed SSU rotation, further head domain swivel (∼12-17°) and nearly complete tRNA translocation relative to the LSU (6, 9). Motions in the EF-G(GDP)-bound ribosome give way to INT3, where head swivel is likely maximal (∼20-26°) and tRNAs are fully translocated with respect to the SSU body (15, 20, 21). Translocation is finalized by full reversal of SSU rotation and head domain swivel, together with EF-G(GDP) release from the POST complex (12, 14, 15). Along with swivel, other orthogonal head domain motions have been associated with translocation, such as tilting away from the subunit interface (20, 22–25). As this framework suggests, the conformational trajectory of translocation is complex, involving multiple degrees of freedom of various molecules and ribosomal domains.

In this study, we sought to use directed evolution and molecular dynamics to explore the flexibility of the conformational trajectory of ribosome translocation. To do this, we focused on the ribosomal RNA (rRNA) ‘hinge’ elements controlling the head swivel motion as targets of mutagenesis and *in vitro* selection. Computational and structural studies have established that the head domain swivels about two 16S rRNA elements within the SSU (26) (**Fig. 2a**). One of these ‘hinges’ (hinge 1) centers around a weak point in helix h28, which is initially kinked and straightens during forward swivel. The other hinge (hinge 2) is a single strand between h34 and h35, moving orthogonally to hinge 1 throughout translocation (**Fig. 2a, b**). Across bacterial and archaeal forms, hinge 1 is substantially more conserved than hinge 2 (27). To explore the sequence and conformational space of hinge function, the hinge elements were mutated and functional variants were selected using the RISE (Ribosome Synthesis and Evolution) platform (28). In this ribosome evolution platform, a ribosomal DNA (rDNA) library is synthesized in ribosome-free cell extract into a ribosome variant pool that is selected for function over multiple rounds (**Fig. 2c**). This strategy yielded multiple alternate hinges capable of translocating the mRNA-[tRNA]_2_ substrate both *in vitro* and *in vivo*, suggesting substantial sequence flexibility in head domain hinge function.

**Figure 2.**
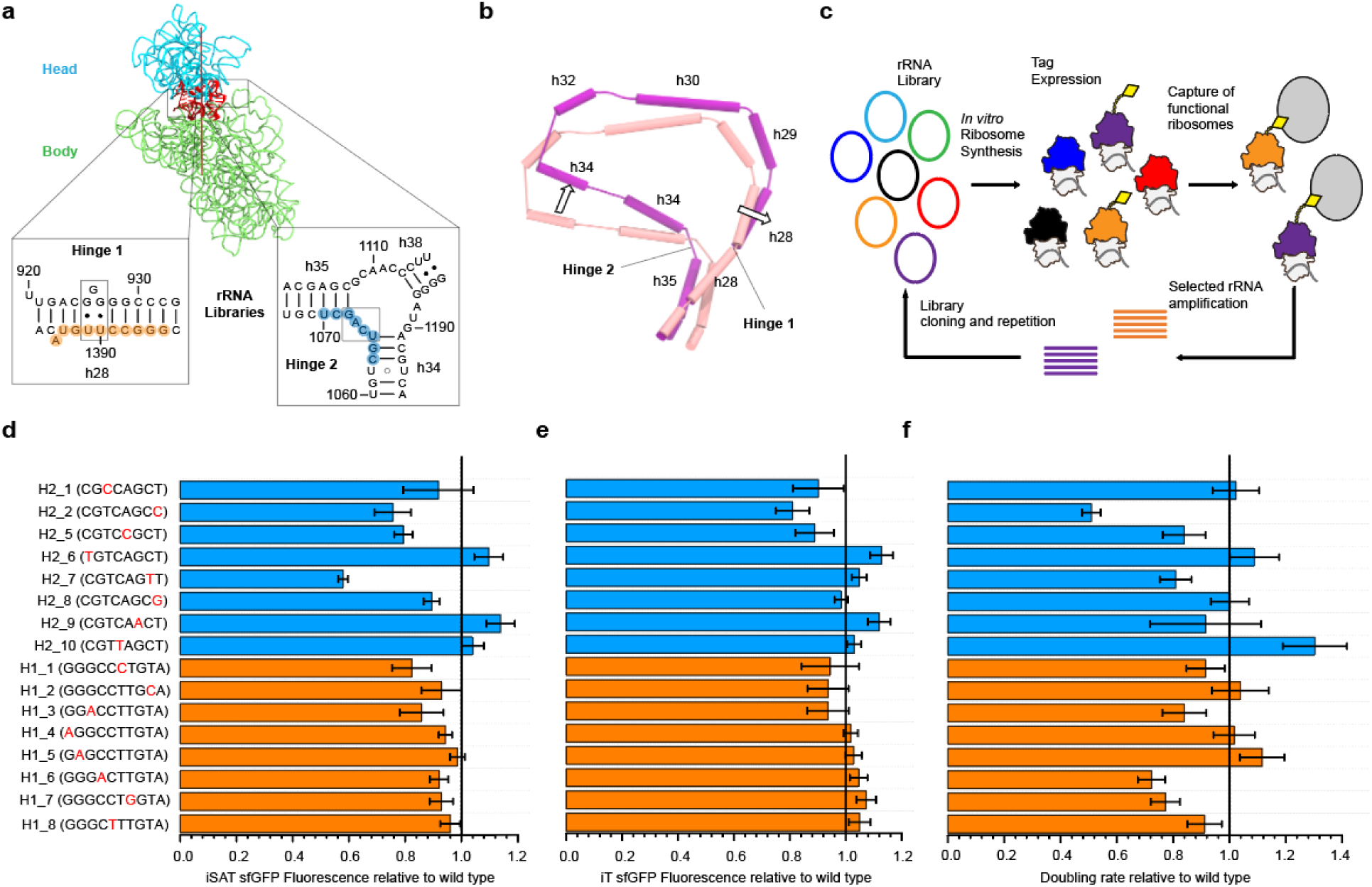
Head domain hinge variants directed evolution and activity. (**a**) Solvent view of the 16S rRNA, showing the head (cyan) and body (green) domains. During translocation the head domain swivels about two rRNA hinge structures. Hinge 1 is a weak point in h28 and hinge 2 is a single-stranded linker between helices h34 and h35. Highlighted in orange and blue are the bases randomized in two separate libraries for hinge 1 and hinge 2, respectively, (PDB ID 4V7B, Reference 9). (**b**) Head swivel and hinge articulation. During swivel hinge 1 motion results from the kinked h28 being straightened, while the orthogonal hinge 2 motion results from h34 swiveling about h35. Cylinders show helical axes in the 16S rRNA core for the PRE (tan) and INT2, swiveled (magenta) conformations. (**c**) Directed evolution was carried out using the RISE (Ribosome Synthesis and Evolution) platform. RISE combines an *in vitro* integrated synthesis assembly and translation (iSAT) system with ribosome display. The platform consists of a library of rRNA variants which are synthesized into ribosome variants in (ribosome-free) S150 cell extract complemented with total ribosome proteins. Functional ribosomes are selected using an expressed tag, and the selected rRNA is amplified and cloned, leading to a new cycle of selection or sequencing. (**d**) Variant activities *in vitro* quantified as the fluorescence of sfGFP synthesized in iSAT reactions relative to wild type. Hinge 1 mutants shown in orange, hinge 2 mutants shown in blue (H2_3 and H2_4 could not be cloned). (**e**) Variant activities *in vitro* quantified as the fluorescence of sfGFP synthesized in iT reactions (translation with prior ribosome synthesis) relative to wild type. (**f**) Variant activities *in vivo*, quantified as the doubling rate of the transformed Squires strains (expressing only the mutant ribosomes) relative to wild type. Wild type cells had a doubling rate of (0.85 ± 0.05) doublings / hour. Error bars correspond to the standard deviation from three independent experiments. The black line denotes wild type activity (=1)..

To explore the potential variability in the translocation conformational rearrangement trajectories dictated by these mutant hinges, all non-hydrogen atom structure-based simulations were performed on EF-G-bound models of the ribosome (20, 23, 29). Analysis of these simulations showed that the hinge points for the head swivel are preserved for the top variants obtained from RISE. However, during the forward swivel, all hinge variants produced smaller fluctuations of the hinge 1 region, suggesting lower flexibility relative to the wild type is selected through hinge mutations. Forward swivel, completed by INT3 formation, was also carried out faster in the selected variants relative to the wild type ribosome, suggesting that delays in achieving the INT3 configuration may not be viable. Interestingly, the three-dimensional head domain trajectory, characterized by head swivel and tilt, was diverse across the variants during the forward and reverse phases of swivel. Similarly, the initial phase of deacyl tRNA motion, from PRE to INT2 formation was notably diverse across variants. These results suggest that a central process in protein synthesis intrinsic to ribosome dynamics, has substantial conformational and kinetic flexibility. Understanding these alternate trajectories can allow us to identify the features of translocation that are critical or non-essential, increasing our knowledge of protein synthesis function, evolution, and engineering.

## Results

### Variant evolution reveals hinge sequence flexibility

To investigate the conformational malleability of mRNA-[tRNA]_2_ translocation specifically, and of protein synthesis in general, we sought to evolve head domain hinge variants. Recent computational and structural studies (26) localized the axis of head swivel between helix h28, the single covalent connection between the head and body domains, and the coaxially stacked helices h35 and h36, which form a non-covalent connection between these domains (**Fig. 2a, b**). The two hinge points of the head domain about this swivel axis were localized inside h28 (hinge 1) and in the single-stranded linker connecting h34 and h35 (hinge 2). With these hinges identified as critical control elements of translocation, selection of alternative hinge structures that permitted protein synthesis was performed. Two separate libraries were constructed, consisting of degenerate mutations in the general region of each head domain hinge, spanning 16S rRNA positions 1385-1394 (Hinge 1 library) and 1063-1070 (Hinge 2 library) (**Fig. 2a**).

Directed evolution of these libraries was carried out *in vitro* to bypass the constraints of cell viability, strain competition and library transformation, focusing solely on the synthetic capacity of the ribosome. A cell-free ribosome synthesis and evolution method called RISE (Ribosome Synthesis and Evolution) (28), was employed to perform this selection (**Fig. 2c**). RISE combines an *in vitro* integrated synthesis assembly and translation (iSAT) system (30) with ribosome display. In this method, a library of rRNA sequence variants (encoded into a pT7rrnB plasmid template) is transcribed and assembled into a ribosome library in (ribosome-free) S150 extract supplemented with total ribosome proteins (TP70). Following translation of a selective peptide (3xFLAG) by functional ribosomes, the ribosome stalls on a stop-less truncated mRNA, and the stalled mRNA-ribosome-peptide complexes are captured using a magnetic resin. The selected rRNA is then purified, reverse transcribed, and either sequenced or assembled into a new library for further RISE cycles.

Previous *in vitro* ribosome synthesis and evolution approaches have carried out mutation mapping of the 70S ribosome (31), enabled assessment of computationally designed ribosomes (32), and evolved the ribosome for new function (33). Here, we applied a modified version of RISE to the two constructed hinge libraries, performing 3 rounds of selection for each (see **Methods**, *Ribosome synthesis and evolution*). Characterized by iSAT sfGFP translation, most of the libraries activity increase was achieved after a single round of RISE (**Table S1**), with minimal improvements from subsequent rounds.

After completing the *in vitro* selection, eight of the top genotypes for each library (ranked by read enrichment after selection relative to the original library, see **Methods**, *Deep sequencing of RISE cDNA*, **Table S2**) were individually cloned and tested in iSAT by translation of sfGFP (**Fig. 2d**). Surprisingly, most of the variants displayed a level of activity close to wild type (> 70%), with no observed correspondence with enrichment rank. Clearly, this measure is indicative of the efficiency of ribosome synthesis and reporter translation, not solely of translocation. Carrying out ribosome synthesis prior to reporter translation (*in vitro* translation or iT, see **Methods**, *iSAT and iT reactions*) generated efficiencies more uniformly comparable to the wild type system (**Fig. 2e**). This suggests that elongation processes (including translocation) have a similar efficiency to wild type ribosomes across the top hinge mutants, and most of the variation observed through iSAT (as in **Fig. 2d**) is likely to stem from ribosome assembly.

To assess the viability of selected variants *in vivo*, the top genotypes were cloned in the ampicillin-resistant pAM552G backbone, containing the rrnB operon under the temperature-sensitive phage lambda (pL) promoter. The constructs were then transformed into the Squires strain (34) SQ171fg, lacking chromosomal rDNA and carrying a pCSacB plasmid with the RiboT-v2 sequence (35). The sucrose sensitivity of pCSacB allows the native tethered ribosome to be replaced by the specific pAM552G ribosome mutant plasmid, and total RNA was isolated from the strains and amplified to confirm the presence of the expected mutations in the 16S rRNA (see **Methods**, *Plasmid replacement and selection in SQ171fg cells*). Doubling rates were derived from the transformed variant strains and compared to wild type (**Fig. 2f**). The mutant cells had varied growth, as demonstrated by doubling rates ranging from ∼50% to 130% of wild type. As with *in vitro* activity, no relationship between enrichment rank and doubling rate was apparent. These rates followed the global trends of the *in vitro* activity, highlighting the reliability of iSAT as an accurate readout of protein synthesis, despite drastically differing conditions as compared to living cells. The viability of most of these strains (15 of the 16 variants have growth rate greater than 70% of the wild type) indicates the capacity of the alternate head domain hinges to carry out protein synthesis in general, and translocation specifically, in the multiple contexts necessary to produce a working bacterial proteome.

### Molecular dynamics simulations of forward and reverse head swivel

To explore the mechanics of translocation for the hinge variant ribosomes, we performed all non-hydrogen atom structure-based simulations of EF-G-bound ribosomes. Models for the various translocation states were constructed using PDB ID: 4V9D (PRE-Classical) (36), PDB ID: 4V7B (INT2) (9), modeled hyper-swiveled state (INT3) (20), and PDB-ID:4V5F (POST) (11) as starting structures. The energy of each model was minimized, and the resulting models were used as either a starting structure or energy minima in translocation simulations. Forward and reverse head-swivel motions of the ribosome were simulated as previously described (20, 23). Forward swivel structure-based simulations were initiated from the PRE conformation of the ribosome with tRNA in the A/P and P/E positions. Stabilizing contacts from the INT3 ribosome conformation were included to define INT3 as an energetic minimum in the simulation. Using this approach, the ribosome head and tRNA were allowed to adopt different configurational space until contacts of the INT3 conformation were adopted. Similarly, reverse swivel simulations, capturing structural dynamics required for the head domain to return to its classical position (POST state) were performed. Here, simulations were initiated in the INT3 state with a head-swivel of ∼26° and were allowed to sample configurational space until the POST state was achieved and stabilized by setting the corresponding contacts as a dominant energy minimum. Reverse head-swivel simulations were performed in the absence of deacylated tRNA to reflect the potential of dissociation of tRNA from the E-site once the INT3 state has been adopted (20).

### Variant core motion preserves hinge points

To investigate if hinge variants introduced structural derivations of the hinges governing head swivel motions, their helical axis were deduced. Specifically, localization of the head domain hinges was performed by measuring position differences in the helical axis of the 16S rRNA between classical and swiveled ribosomes (26). A series of helices leads from the ribosome body and forms the head domain. On one end, h28 (the ‘neck’) makes a covalent connection from the head to the body domain of the SSU, where it stacks coaxially with h1, h2 and h3. At the other end, h36 forms a non-covalent bridge with h2 via A-minor interactions. The core helices h28-h29-h30-h32-h34-h35-h36 lead from and to the body, forming the head domain (**Fig. 3a**). Based on these helices, a continuous helical axis for the head has been calculated, and the motion across the head due to swivel can be quantified (26) (**Methods**, *Calculation of core helical axis motion*). The displacement of the helical axis between the simulated (16° swivel) swiveled structures relative to the classical configuration was calculated for the wild type and top variants. This quantification showed that the wild type hinge points are preserved for all variants, localizing to inflection points near 16S rRNA positions 928 and 1065 (**Fig. 3b, c**). The hinge points were preserved for the reverse swivel motion as well (**Figs. S1c, d**). Intriguingly, we observed a broad distribution of helical axis displacements, deviating over 5 Å in some trajectories relative to wild type ribosomes.

**Figure 3.**
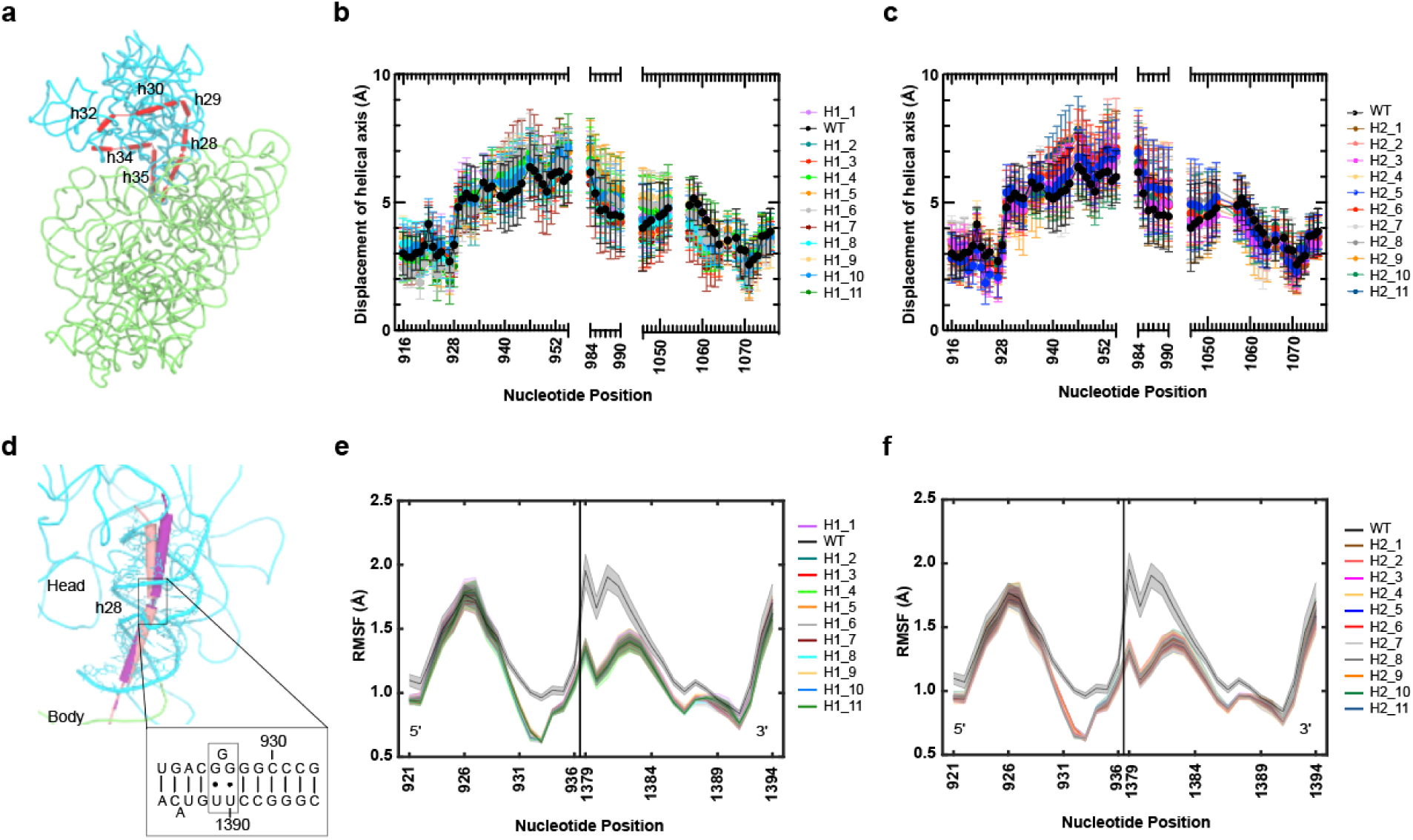
Variant helical axis and hinge motions. (**a**) Calculated helical axis (red) for the structural core of the 30S head in 16S rRNA. The head (domain III) is shown in cyan, and the body (domains I, II and IV) is shown in green (PDB ID 4V7B, Reference 9). (**b**) Displacement at each position along the calculated helical axis from the classical PRE state to the simulated conformation at 16° swivel for wild type and top selected hinge 1 variants. (**c**) Same as (b) for wild type and top selected hinge 2 variants. (**d**) Hinge 1 motion in the wild type ribosome. Hinge 1 is positioned in a weak point in h28 and its motion results from the kinked helix h28 being straightened during swivel. Cylinders show helical axes in the 16S rRNA core for the classical PRE (tan) and swiveled INT2 (magenta) conformations. (**e**) Root-mean-square-fluctuations (RMSF) of hinge 1 during forward swivel for wild type and top selected hinge 1 variants. (**f**) Same as (e) for wild type and top selected hinge 2 variants.

These distinctions indicate that although the head-swivel angle is maintained, the helices themselves can be arranged in different configurations to achieve these angles.

### Reduced hinge fluctuations allow effective translocation

To quantify the motion of the hinges throughout forward head swivel, the root mean square fluctuation (RMSF) across the hinge regions was measured (**Methods**, *Hinge fluctuations*). Interestingly, hinge 1 fluctuations (**Fig. 3d**) were reduced relative to wild type for all hinge variants (including hinge 2 variants) during forward head swivel (**Figs. 3e, f**). In contrast to these results, hinge 2 fluctuations during forward swivel and hinge 1 and 2 fluctuations during reverse swivel were not significantly altered in the variants (**Fig. S1**). This suggests that reduced flexibility across h28 during forward swivel is a selectable feature and that hinge 1 may exhibit more conformational plasticity during forward swivel motions, whereas fluctuations for reverse swivel are more limited. To understand if the change in fluctuations can be reflected in the global motions of hinge 1, a principal component analysis was performed for hinge 1 forward head-swivel motions. Subtle alterations were observed for the first principal component describing the movement of hinge 1 for representative variants (**Figs. S2a-c**). Projection of these motions onto the structure of hinge 1 indicates that variants undergo motions parallel to the helical axis of hinge 1 (**Figs. S2d-f**). This indicates that the dominant motion of hinge 1 during forward swivel is not altered by its flexibility.

### Variant translocation displays flexible head domain trajectories

Following EF-G binding, the head domain is thought to swivel and tilt to translocate the mRNA-[tRNA]_2_ substrate (7, 15, 20, 23). This swivel-tilt coordination controls the opening of the SSU corridor that the translocation substrate moves through (9, 23). To study the global conformational flexibility of translocation in the hinge variants, multiple degrees of freedom of the head domain were considered. Specifically, the trajectory of the head was described by both the head swivel and the orthogonal head tilt (**Fig. 4a**). Using the head swivel and tilt angles, the global trajectories of the head domain relative to the SSU body were described. To compare the variant trajectories to that of wild type, heat maps showing the probability of conformations during forward swivel were derived. Simulations for the variants were carried out for the steps required for the wild type ribosome to reach maximum swivel and final tRNA positioning (INT3). Notably all the variants studied reached this state faster than wild type (**Fig. 4b**, **Figs. S3, S4**), completing forward swivel (78 ± 5) % faster (**Table S3**).

**Figure 4.**
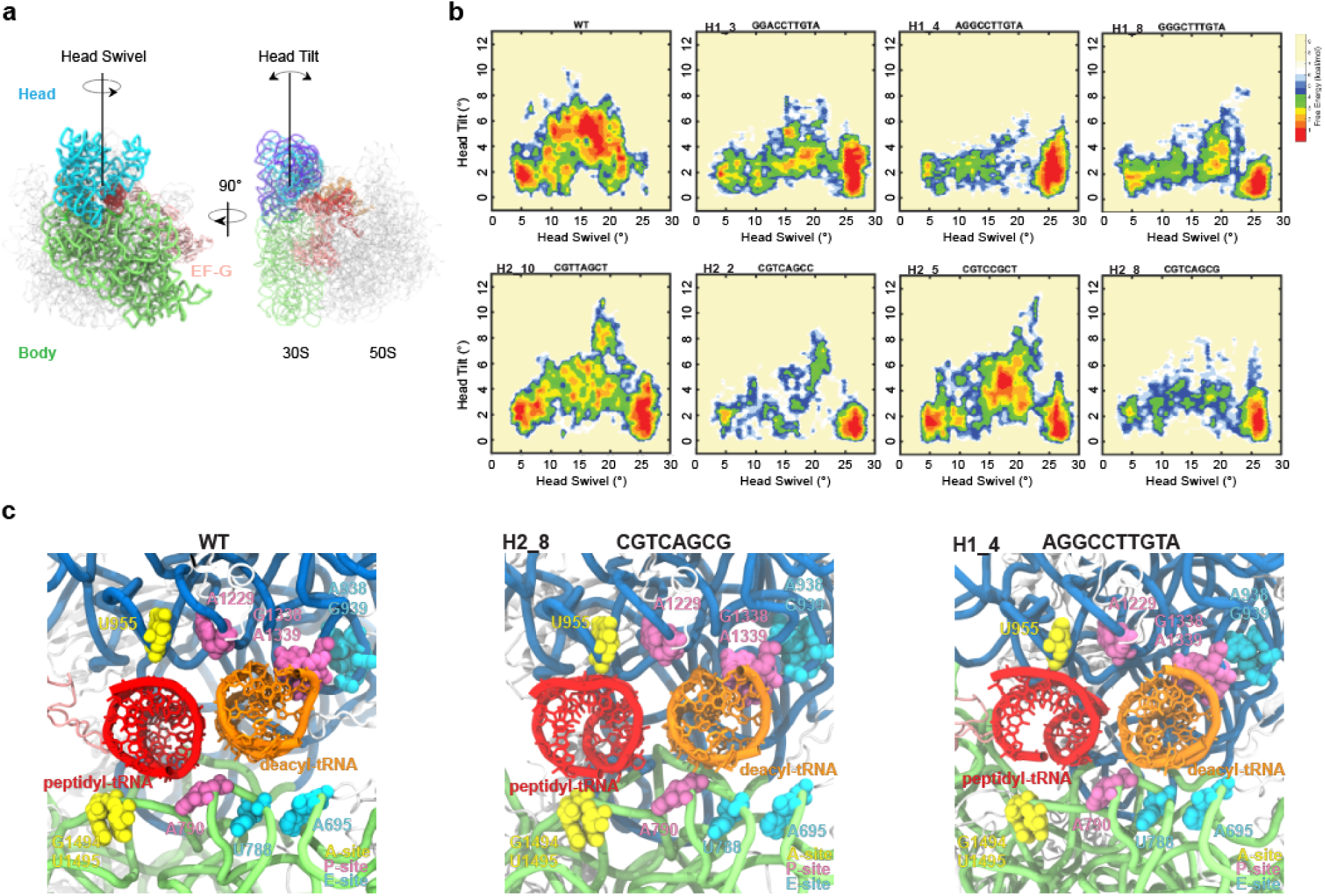
Alternate translocation trajectories of head domain motion. (**a**) The main head domain motions during translocation. Solvent view of the 30S subunit (Left) showing head swivel in the direction of translocation. 70S particle (Right) showing the direction of head tilt, orthogonal to head swivel. (**b**) Free-energy landscapes of the forward swivel (PRE to INT3) stage of translocation with head domain swivel and tilt angles as reaction coordinates. The landscapes are based on Boltzmann weighted probability distributions. The heat map is a normalized number of frames in the targeted MD simulation, where the frame count is divided by the maximum frame count. Shown are the maps for the wild type and seven representative hinge variants active *in vitro* and *in vivo*. Maps correspond to ten trajectories per genotype. (**c**) Interactions between the small subunit and tRNAs for simulated translocation intermediates. Subunit interface view of the simulated wild type and representative variant conformations at 16° of forward swivel showing tRNA contacts in the A (yellow), P (magenta) and E (teal) sites.

The wild type ribosome trajectory of the head domain describes an arc-like tilt throughout the forward swivel. The tilt angle rises from ∼2° at initial (∼0°) head swivel to a maximum tilt of ∼6° as the head swivel nears ∼17°, as observed in previous intermediate structures (6, 37). Tilt then reverts to ∼1° at maximum (∼23°) swivel. The state observable at ∼4°-6° tilt and ∼15°-18° swivel corresponds to the INT2 translocation intermediate (6, 7, 15, 20), displaying a large degree of swivel and tRNA motion and leading to complete substrate repositioning at INT3. Importantly, the selected variants displayed a variety of head domain trajectories. While certain trajectories were closer to the wild type ribosome (**Fig. 4b**, H2_5), some had a substantially reduced head tilt (**Fig. 4b**, H1_4, H2_8) and others showed alternative intermediate conformations (**Fig. 4b**, H1_3, H1_8, H2_2, H2_10).

An important consequence of this variation in the degree of head tilt during forward swivel is the flexible extent of the platform opening and of the small subunit ‘grip’ on the tRNAs. Forward swivel simulation snapshots (at 16° swivel) show the extent of this variability (**Fig. 4c**). As established previously, these structures show that during forward swivel specific head-tRNA contacts are maintained, while body-tRNA contacts are disrupted (6, 8, 9). The different tilt profiles, however, result in variation of the proximity and extent of the 30S-tRNA contacts. As small tilt angles (< 4°) can achieve the fully translocated tRNA positions at INT3, this indicates that the forward tilt motions and opening of the platform is not a requisite feature for tRNA translocation.

In contrast to forward swivel, the reverse swivel for the wild type was completed faster, with transit time being (55 ± 10) % that of the variants (**Table S3**). As in the forward swivel, the wild type ribosome tilt trajectory for the head described an arc, and the selected mutant trajectories were variable. Unlike forward swivel, all the variant reverse swivels consistently achieve significant (∼7°-12°) degrees of tilt (**Figs. S5, 6**). Under this framework, reverse head swivel requires more tilt than forward swivel, but large tilt angles are not essential to achieve the POST position.

### Variant translocation displays flexible tRNA trajectories

To describe the observed conformational flexibility of translocation from the perspective of substrate motion, the trajectory of the incoming deacyl tRNA into the E-site was monitored. This was done by measuring the distance between the deacyl tRNA and the E loop in the SSU body (h23, G691-A695) (**Fig. 5a**). Using this distance and the extent of head swivel, the tRNA trajectory was described in heat maps for wild type and variants (**Fig. 5b, Figs. S7, 8**). The wild type ribosome motion of deacyl tRNA occurs through two main stages. The first stage displays fast tRNA motion, going from an initial distance to the E loop of ∼32 Å (at ∼5° swivel) to ∼23 Å (at ∼10° swivel). This stage takes only (25 ± 3) % of the time required to reach maximum swivel (**Table S3**). The second stage consists of slower tRNA motion from ∼23 Å (at ∼10° swivel) to ∼20 Å (at ∼25° swivel). This latter stage includes the transition from INT2 (∼15° swivel) to INT3. Exchange between these states is thought to consist of an accurate final positioning of the translocation substrate (7, 15, 20).

**Figure 5.**
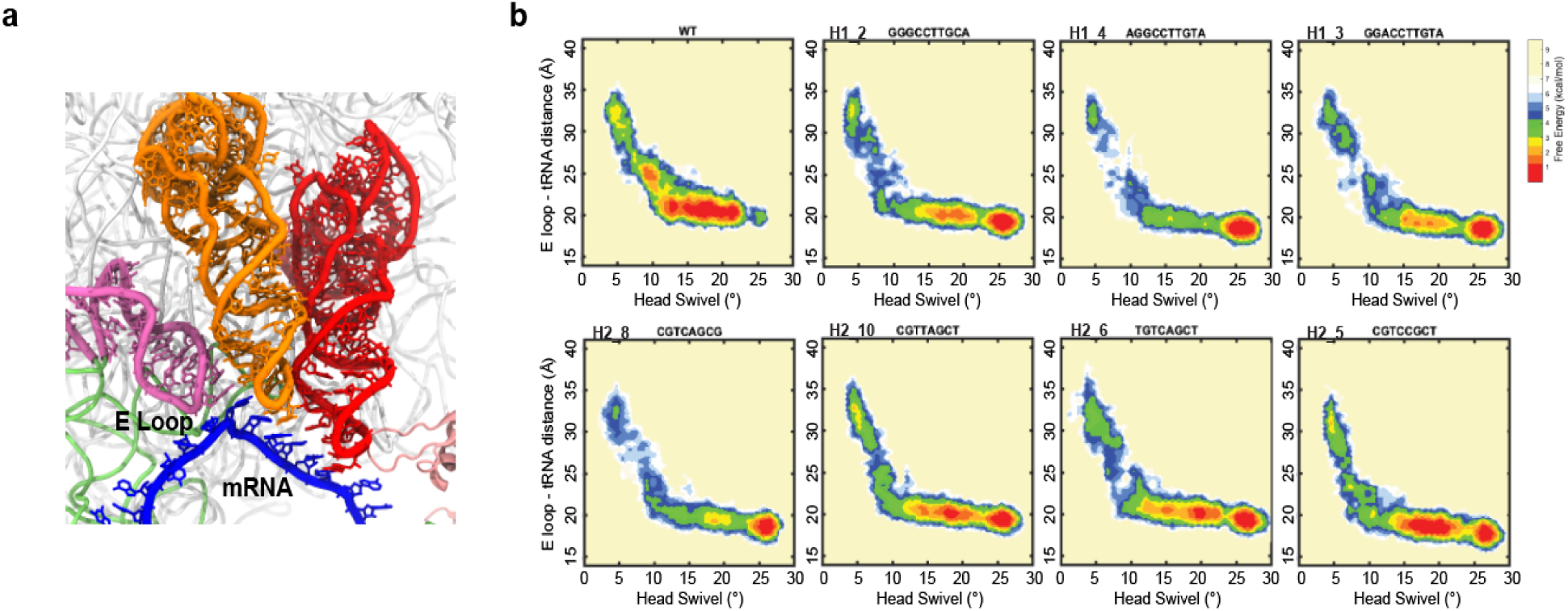
Alternate translocation trajectories of tRNA motion. (**a**) Deacyl tRNA motion during forward swivel was tracked by its distance to the E loop (h23, G691-A695). (**b**) Free-energy landscapes of the forward swivel (PRE to INT3) stage of translocation with head domain swivel angle and E loop-Deacyl tRNA distance as reaction coordinates. The landscapes are based on Boltzmann weighted probabilit distributions. The heat map is a normalized number of frames in the targeted MD simulation, where the frame count is divided by the maximum frame count. Shown are the maps for the wild type and seven representative hinge variants active *in vitro* and *in vivo*. Maps correspond to ten trajectories per genotype.

Interestingly, tRNA motion for the selected mutants varied widely in the initial, fast phase of translocation (**Fig. 5b**). This initial phase displays varying transit of the previously observed, short-lived intermediate INT1 (∼30 Å, ∼6° swivel) (6, 7, 15, 20), with some clear INT1 dwells (H1_2, H1_3, H2_10, H2_6, H2_5) and some nearly absent dwells (H1_4, H2_8). The following tRNA motion phase (∼10° to ∼25° swivel), though still faster than wild type for all variants, is uniformly the slower stage of forward swivel (**Table S3**). Notably, the dynamics of this phase are considerably uniform across variants.

## Discussion

In this study, mutant rRNA head domain elements capable of protein synthesis were obtained through directed evolution. By characterizing translocation in these variants through molecular simulations, alternate conformational trajectories for ribosome translocation were identified. Focusing on the hinge elements controlling head domain swivel, the variants selected using *in vitro* ribosome synthesis and evolution platform provided hinge mutants capable of sustaining ribosome function *in vitro* and *in vivo*, including mutations in nucleotides considered to be universally conserved (27) (H1_7). These variants, and their uniform capacity to carry out translation in iSAT and in living cells (**Fig. 2d-f**), suggest that there is substantial sequence space available to the hinges to perform translation well enough to support life.

Though the translocation simulations demonstrate similar hinge points across the selected mutants (**Figs. 3b, c**, **Figs. S1c, d**), the fluctuations of hinge 1 throughout forward swivel were notably diminished for all variants (**Figs. 3e, f**). This, together with variant increased forward swivel rate (**Table S3**), suggests that the variant hinges produce a more concerted motion towards the maximum swivel state relative to wild type ribosome. An apparent consequence of this concerted hinge variant movement during forward swivel is an average head tilt that is similar to or reduced relative to the wild type ribosome (**Fig. 4b**, **Fig. S3**). This result implies that forward head domain motions do not rely upon a specific tilt trajectory to effectively open the SSU corridor, and that a unique profile of 30S-tRNA contacts is not necessary for translocation (**Fig. 4c**). The observation that multiple different head tilt trajectories can translocate the mRNA-[tRNA]_2_ substrate indicates a high degree of conformational flexibility in translocation.

Though variant reverse swivel is conformationally diverse and transpires slower than for wild type (**Table S3**) a minimal, substantial degree of tilt (≳ 7 °) is uniformly observed (**Figs. S5, S6**), suggesting a steric constraint for reverse head swivel. As deacylated tRNA in the E-site is considered to disengage from the ribosome at the INT3 state (20), it was not included in the simulations, indicating that this minimum tilt does not arise from steric blocks imposed by the E-site tRNA. The other confounding steric interactions of the head domain leading to this tilt could be with the peptidyl tRNA through the AP loop (nucleotides 954-957 of 16S rRNA) of the head domain (**Fig. 4c**). This steric barrier may be a contributing factor to the >98% conservation of this region in ribosomes, excluding mitochondrial ribosomes (20).

Further alternate features of translocation were found in the variant deacyl-tRNA motion. For the initial PRE to INT2 substrate motion, the intermediate profiles are diverse between wild type and variant ribosomes (**Fig. 5b**, **Figs. S7, S8**). This implies that state INT1 (described in previous studies (7, 15, 19)) transit can vary substantially while yielding efficient and accurate translocation *in vitro* and *in vivo*. In contrast, other motions critical to translocation are substantially conserved, notably INT2 and INT3 transit during the second, slower phase of deacyl tRNA motion, or the balance between head swivel and small subunit rotation (**Figs. S9, S10**).

Our results suggest features of translocation, such as the head tilt magnitude and sampling of stable intermediates, are contingent to the global function of protein synthesis in general and to translocation in particular. If not necessary, how can the prevalence of these aspects of ribosome function be explained? One possibility is that these specific features are not functionally necessary for forward swivel but arise from degrees of freedom involved in other processes. For example, substantial head tilt seems to be non-essential during forward swivel yet is required in reverse swivel (**Figs. S5, 6**), as well as in trans-translation (38) and in translating ‘slippery’ proline sequences (39). Another option is that the specific conformational trajectory of wild type translocation was necessary at a previous point of ribosome evolution and subsequent alterations have rendered it malleable (40). A candidate evolutionary shift for this hypothesis is the development of elongation factors, notably EF-G (41). Prior to the evolution of these factors, the ribosome may have translated proteins as a standalone enzyme. To achieve translation without auxiliary factors it may have had to adopt additional degrees of freedom or stable intermediates to overcome energetic barriers. These energetic barriers could be alleviated in the presence of extant auxiliary factors, such as EF-G, during translocation and translation. Finally, it is plausible that these features of wild type translocation are not necessary for protein synthesis or cellular growth under the conditions studied, but concede advantages under different conditions or increased adaptability (42). This could include starvation conditions where translation speeds are altered (43) or temperature fluctuations (44), where additional degrees of freedom may facilitate translocation.

This study highlights the flexibility of the conformational pathways of the ribosome during translocation. In doing so, we have established some of the critical degrees of freedom and intermediate states of this process while indicating motions that are more variable or contingent. This understanding is of importance in the engineering of the translation machinery, and in the targeting of translation with potential therapeutics or antibiotics, as perturbing essential processes will yield more robust treatments.

## Methods

### Plasmid and library construction

The plasmids pT7rrnB (containing the rDNA operon rrnB under a T7 promoter), pJL1sfGFP (encoding the reporter sfGFP under a T7 promoter, Addgene accession #102634) and pRDV3xFLAG-TolA-HH (encoding the 3xFLAG tag for ribosome selection under a T7 promoter) were used in *in vitro* integrated synthesis assembly and translation (iSAT) reactions (30, 45, 46). The plasmid backbone pAM552G (35, 47) (Addgene accession #154131) was used for testing ribosome variants from RISE *in vivo* (see below). pAM552G carries Ampicillin resistance and a copy of the rrnB operon under the temperature-sensitive phage lambda (pL) promoter. For library creation and cloning, a pT7rrnB variant with a 438 bp deletion in the 16S gene (pT7rrnBΔ438) was created by inverse PCR, including unique cut sites for the off-site type IIs restriction enzyme SapI. Libraries of rDNA were generated by cloning PCR amplified (with SapI sites) library amplicons ordered from Epoch Life Science and ligating them into the SapI-digested pT7rrnBΔ438 variant. The reconstituted library was then transformed into MegaX DH10B T1 R electrocompetent cells (ThermoFisher), cultured and purified for the initial RISE cycle. The top rDNA variants enriched from RISE were constructed on pT7rrnB and pAM552G backbones for *in vitro* and *in vivo* testing, respectively. The variants were created from the wild type plasmids using the Q5 Site-Directed Mutagenesis Kit (NEB) and transformed into MegaX cells made chemically competent (Mix & Go *E. Coli* Transformation Kit, Zymo Research). The top 8 variants of each library selection were constructed, though H2_3 and H2_4 could not be cloned. pAM552G variants were incubated at 30°C to prevent plasmid rRNA expression. Full plasmid nanopore sequencing was carried out to confirm the expected variant genotypes.

### iSAT and iT reactions

iSAT reactions were performed by mixing a master mix containing salts, substrates, rDNA plasmids/libraries, reporter/tag plasmids, and cofactors with ribosome-free MRE600 S150 extract and total proteins of 70S ribosomes, TP70 (28). The final concentrations of the main additions were ∼2 mg/mL for the S150 protein content, 100 nM TP70, 1.5 μM T7 RNA polymerase, 2.2 nM (11 ng/μL) rDNA encoding plasmids or 0.36 nM (1.8 ng/μL) rDNA libraries, and 2.6 nM (3.9 ng/μL) pJL1sfGFP, pRDV 3xFLAG. Reactions were mixed and incubated at 37°C for 3 hours (for sfGFP expression) or 90 minutes (for RISE). To monitor protein synthesis alone from pre-assembled ribosomes (*in vitro* translation or iT), reactions with no reporter plasmid were incubated at 37°C for 1h to mature the ribosomes, followed by addition of the pJL1sfGFP plasmid. sfGFP expression was monitored using a Bio Rad CFX Touch Real-Time PCR Detection System with a 96 well plate reader.

### Ribosome synthesis and evolution

RISE selections were performed as in a previous study (28) with some modifications. iSAT reactions of 12 μL were carried out with the different rDNA libraries and pRDV 3xFLAG as a ribosome display tag. The anti-ssRA oligonucleotide (5’-TTA AGC TGC TAA AGC GTA GTT TTC GTC GTT TGC GAC TA−3’) was included at 5 μM to hinder stalled ribosome dissociation. Reactions were incubated for 90 minutes at 37°C. Following incubation, reactions were kept at room temperature throughout the protocol and diluted with 15 volumes (180 μL) of wash buffer with Tween^®^, WBT (50 mM Tris-acetate (pH 7.5 at 4°C), 50 mM magnesium acetate, 150 mM NaCl and 0.05% Tween^®^ 20). For each RISE reaction 10 μL packed gel volume of ANTI-FLAG M2 magnetic beads (Sigma-Aldrich) were washed three times with 200 μL wash buffer, WB (50 mM Tris-acetate (pH 7.5 at 4°C), 50 mM magnesium acetate and 150 mM NaCl). The diluted iSAT reactions were added to the washed beads and incubated with gentle rotation for 20 minutes. Reactions were then washed 5 times with WBT buffer, 2 times with WB buffer and resuspended in elution buffer, EB (50 mM Tris-acetate (pH 7.5 at 4°C), 150 mM NaCl and 50 mM EDTA (Ambion)), incubating the suspension with gentle rotation for 30 minutes. The supernatant was recovered from the beads and the rRNA was purified using a Qiagen RNeasy MinElute Cleanup Kit. RT-PCR of this rRNA was done using SuperScript III One-Step RT-PCR System with Platinum *Taq* DNA Polymerase (Invitrogen) in 100 μL reactions according to product literature. The RT-PCR was performed with primers that amplified the bases 1026–1476 of the 16S rRNA gene and included SapI cut sites. This amplicon was then used for next-generation sequencing or cloned into the pT7rrnBΔ438 variant and amplified for further RISE cycles.

### Deep sequencing of RISE cDNA

Libraries with one variable region were sequenced with 250 bp paired-end, non-overlapping Illumina NGS at the University of Minnesota Genomics Center. Since the regions of interest for each of the libraries studied (Hinges 1 and 2) are located close to the ends of the primer regions, no alignment was necessary. Sequences were extracted from the raw reads using Python, searching for up- and downstream sequences of N+2 (e.g. for a variable region of 8 nt, 10 nt up stream and 10 nt downstream were queried). No mismatches were allowed in the search sequences. Sequences containing nucleotides with phred-score of less than 30 (p>0.001) were discarded. Extracted sequences were trimmed and counted for both the original input library and the sequence pool after 3 rounds of RISE selection. Variant abundance was calculated as counts of each sequence relative to all sequence counts of the library. Enrichment was calculated as variant abundance in output library relative to abundance in the original library. Winners were determined via ranking by highest enrichment.

### Plasmid replacement and selection in SQ171fg cells

Electrocompetent *E. coli* SQ171fg cells containing a pCSacB plasmid (granting sucrose sensitivity) with kanamycin resistance (KanR) were prepared. The pCSacB/KanR plasmid carries the sequence for the engineered, subunit-tethered ribosome Ribo-Tv2 (35), serving as the only rRNA operon in the cell. Variant ribosomes of interest in the ampicillin-resistant pAM552G backbone can be transformed into the cells, and the original pCSacB-RiboTv2-KanR plasmid can be replaced by culture with sucrose and carbenicillin. This replacement and cell viability based on the variant ribosomes is confirmed by susceptibility to Kan (32). 150 to 300 ng of each variant plasmid in the pAM552G backbone was transformed into 50 μL of electrocompetent SQ171fg cells. Cells were recovered in 800 μl of SOC medium in 1.5 ml tubes at 37°C with shaking at 250 rpm for one hour. After recovery, 1.2 ml of SOC with 0.42% sucrose and 100 μg/mL were added to the cells (60 μg/ml carb, 0.25% sucrose final concentrations). These 2 ml cultures were further grown in 50 mL falcon tubes at 37°C overnight. The overnight cultures were pelleted, resuspended in 150 μL LB medium and plated in LB agar plates with 5% sucrose and 100 μg/mL carb.

The plates were incubated until colonies were clearly visible (24-72 hours). If no colonies were observed for a specific variant, the transformation was repeated twice more. Colonies were picked from each plate and grown overnight in LB medium with Carb_100_. These cultures were then preserved as glycerol stocks and spotted (3 μL) in both Carb_100_ and Kan_50_ plates. Colonies that grew in carb plates only were grown from glycerol stocks in LB with Carb_100_ and further analyzed. Total RNA was isolated from these cultures using Ambion Trizol^TM^ LS. RT-PCRs of this total RNA were performed amplifying a 635nt region of the 16S rRNA containing both hinge library sites. These amplicons were then nanopore sequenced, and the presence of the expected mutations was confirmed. For all hinge variants capable of supporting life, cell doubling rates were measured. To do this, overnight cultures were grown in LB with Carb_100_ from glycerol stocks. We inoculated 200 μL of LB with 5 μL of each overnight culture and recorded the OD_600_ every 5 minutes over 5 hours with 15-second mixing in between reads in a SPECTRAmax^®^ 340PC Microplate Spectrophotometer with a qPCR seal.

### Model generation for molecular simulations

Molecular simulations were performed using previously derived models of the PRE, INT3, and POST *E. coli* ribosome conformations (20). The PRE conformation was modeled from PDB ID: 4V9D (36) with tRNA in the A/P and P/E positions. The INT3 conformation model was generated previously (20). In brief, the INT3 conformation was modeled by restraining the coordinates of the SSU body from PDB ID: 4V7B and rotating the head domain to a 22.6° head swivel angle along the same helical axis as observed between the INT1 and INT2 states (9). RMSD restraints were used to treat the SSU body, EF-G, LSU, L1, tRNA, and mRNA as rigid bodies to achieve the INT3 conformation. NAMD 2.12 (48) with CHARMM C36 force fields (49) was implemented to minimize and equilibrate the INT3 conformation. Lastly, the POST conformation was modeled from PDB ID: 4V5F (11) with tRNA in the P/P and E/E positions. The mRNA bound to each ribosome model was converted to 5’-GGAAAAAUGUUUAAAAAA-3’, encoding for tRNA^fMet^ and tRNA^Phe^.

Hinge variants were introduced into the 16S rRNA using the Swapna package in Chimera (50). To each model a water box of SPC/E model waters were added extending 15 Å beyond the extremities of the ribosome in addition to KCl and MgCl_2_ ions to a concentration of 30 mM and 7 mM, respectively with GROMACS 2022 (51, 52). Each hinge variant was minimized in GROMACS 2022 using an iterative steepest descent approach by minimizing the potential energy of the water proceeded by the ribosome sidechains and backbone.

### All-atom structure-based simulations

All-atom structure-based simulations were performed using GROMACS 2022 software with forcefields generated by the SMOG-2.4.2 software (53). Simulations were performed at a temperature of 0.5ε, where ε is reduced units. The functional form of the potential used in the simulations is defined by:

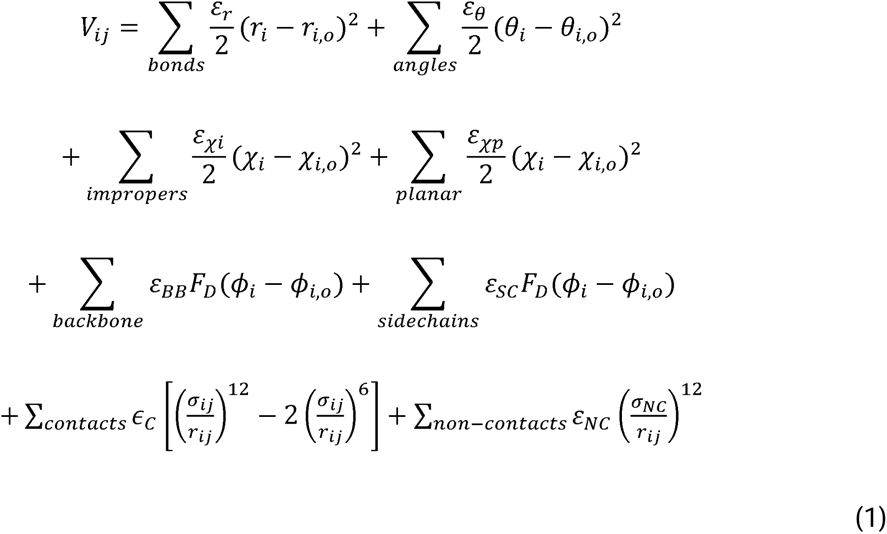

where *ε_r_* = 50 *ε*_0_, *ε_θ_* = 40 *ε*_0_, *ε_Xi_* = 10 *ε*_0_, *ε*_Xp_ = 40*ε*_0_, *ε_NC_*= 0.1 *ε*_0_, *σ_NC_* = 2.5 Å, and *ε*_0_= 1 and 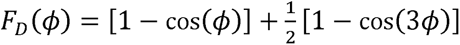. Contacts were determined using a cutoff distance of 4 Å between heavy atoms. The strength of the contributions between the mRNA and tRNA substrate with the ribosome were rescaled by 0.6 to allow for translocation of the substrate. Additionally, the interface contacts between the SSU head and body were scaled by 0.3 to allow for head swivel motions. Contacts between the SSU and LSU were scaled by 0.3 for intersubunit contacts within 20 Å of core and by 0.1 for those outside this radius. This contact scaling approach has been utilized to accurately simulate translocation from the PRE to POST conformation of the ribosome previously (23). Finally, interactions involving EF-G with ribosomes or tRNA were adjusted by a factor of 0.6 to account for the temporary nature of EF-G binding. Simulations were performed for 1.5 x 10^7^ timesteps with a step size of 0.002, as previous simulations of this type have indicated that 1 reduced time unit (t_ru_) is ∼1ns (54, 55), each simulation is ∼30 μs. At minimum 10 simulations were performed for wild type ribosomes and each variant, totaling ∼6.9 ms of simulations time.

### Calculation of core helical axis motion

The helical axis of SSU head swivel motions were determined using the Curves+ software (56, 57). Specifically, the helical axis of h28, h29, h30, h32, h34, h35, and h36 were determined using snapshots of molecular simulation trajectories of ribosomes when they had reached the PRE (0°), INT2 (16°), INT3 (22.6°), or POST (0°) states defined by specific head swivel angles. Additionally, an intermediate in the transition between INT3 and POST was used to investigate the helical axis of reverse head-swivel at an angle of 13°. This angle was chosen as an intermediate from free-energy landscape approximations.

### Hinge fluctuations

Backbone root mean squared fluctuation (RMSF) of hinges was measured using GROMACS 2022 (51, 52). RMSF was calculated with:

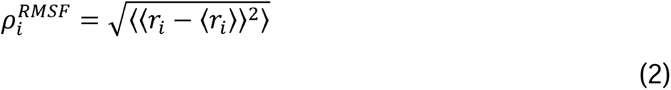

where *r_i_* are the coordinates of atom *i* and 〈*r_i_*〉 are the time averaged coordinates.

### Head swivel and tilt measurements

Head swivel and tilt measurements were performed as previously described (20, 23). In brief, a set of core residues were utilized to align residues of the 16S body to a structural reference PRE configuration and the vector of the reference is measured from a plane derived from three fixed atoms of A977 and A1374 P and U1298 O2P. Subsequently core residues of the head domain are used to align the reference head to those of the trajectory and the vector is measured again from the same vector.

Comparing the vectors from the simulation to the reference allows for quantification of the tilt and head swivel angles.

### Body rotation measurements

Similar for the measurements of head tilt and swivel, rotations of the SSU body were measured using a plane derived from atoms P of C237 and C811 and O6 from G1488. The model from the simulations in each frame was aligned to a reference using core residues in the 23S, then the vector for the plane is measured. Subsequently, the SSU body reference structure is aligned to the model from the simulation using core residues of the 16S from the body and the vector of the plane is determined. The angle between the two vectors is then used to measure the body rotation during translocation.

### tRNA movement measurements

tRNA movement was assessed by measuring the distance between the O3’ atom of G768 of the 16S rRNA and the O3’ atom of A35 of the peptidyl tRNA. Distance measurements were performed in GROMACS 2022. This reaction coordinate was selected to describe the barrier of tRNA movement from the P/E to E/E position during translocation.

## Acknowledgements

We thank Scott Blanchard for thoughtful review of the manuscript. This work was supported by the Alfred P. Sloan Foundation grant G-2022-19420 (K.P.A.), Howard Hughes Medical Institute Hanna Gray fellowship grant GT11084 (J.L.A.), the Arkansas Biosciences Institute (D.G.), the major research component of the Arkansas Tobacco Settlement Act of 2000 (D.G.); and the start-up fund of the University of Arkansas (D.G.). M.C.J. and J.A.W. gratefully acknowledge support from the Army Research Office [W911NF-16-1-0372] and Army Contracting Command [W52P1J-21-9-3023].

## Author contributions

J.L.A. and D.G. designed the experiments and simulations. J.L.A. and M.J.H. designed the libraries used for RISE. J.L.A. and D.G. analyzed the experimental and simulation data. J.A.W. analyzed the deep sequencing data. All authors contributed to the writing of the manuscript.

---

**Figure S1.**
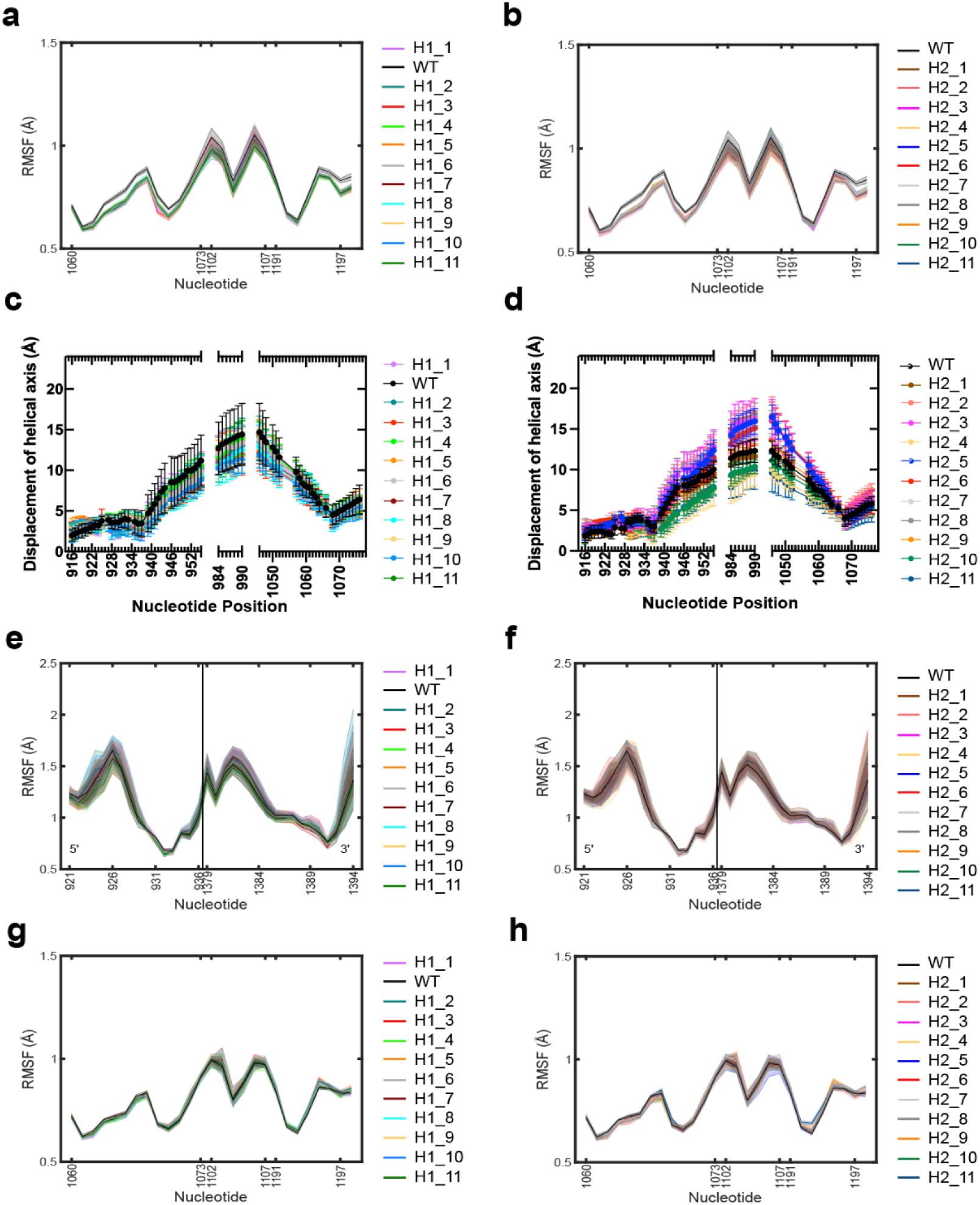
Hinge variant simulation swivel rates relative to wild type. (**a**) Root-mean-square-fluctuations (RMSF) of hinge 2 during forward swivel for wild type and top selected hinge 1 variants. (**b**) Same as (a) for wild type and top selected hinge 2 variants. (**c**) Displacement at each position along the calculated helical axis from the maximum swivel (INT3) state to the intermediate (13° head swivel) conformation for wild type and top selected hinge 1 variants. (**d**) Same as (c) for wild type and top selected hinge 2 variants. (**e**) Root-mean-square-fluctuations (RMSF) of hinge 1 during reverse swivel for wild type and top selected hinge 1 variants. (**f**) Same as (e) for wild type and top selected hinge 2 variants. (**g**) Root-mean-square-fluctuations (RMSF) of hinge 2 during reverse swivel for wild type and top selected hinge 1 variants. (**h**) Same as (g) for wild type and top selected hinge 2 variants.

**Figure S2.**
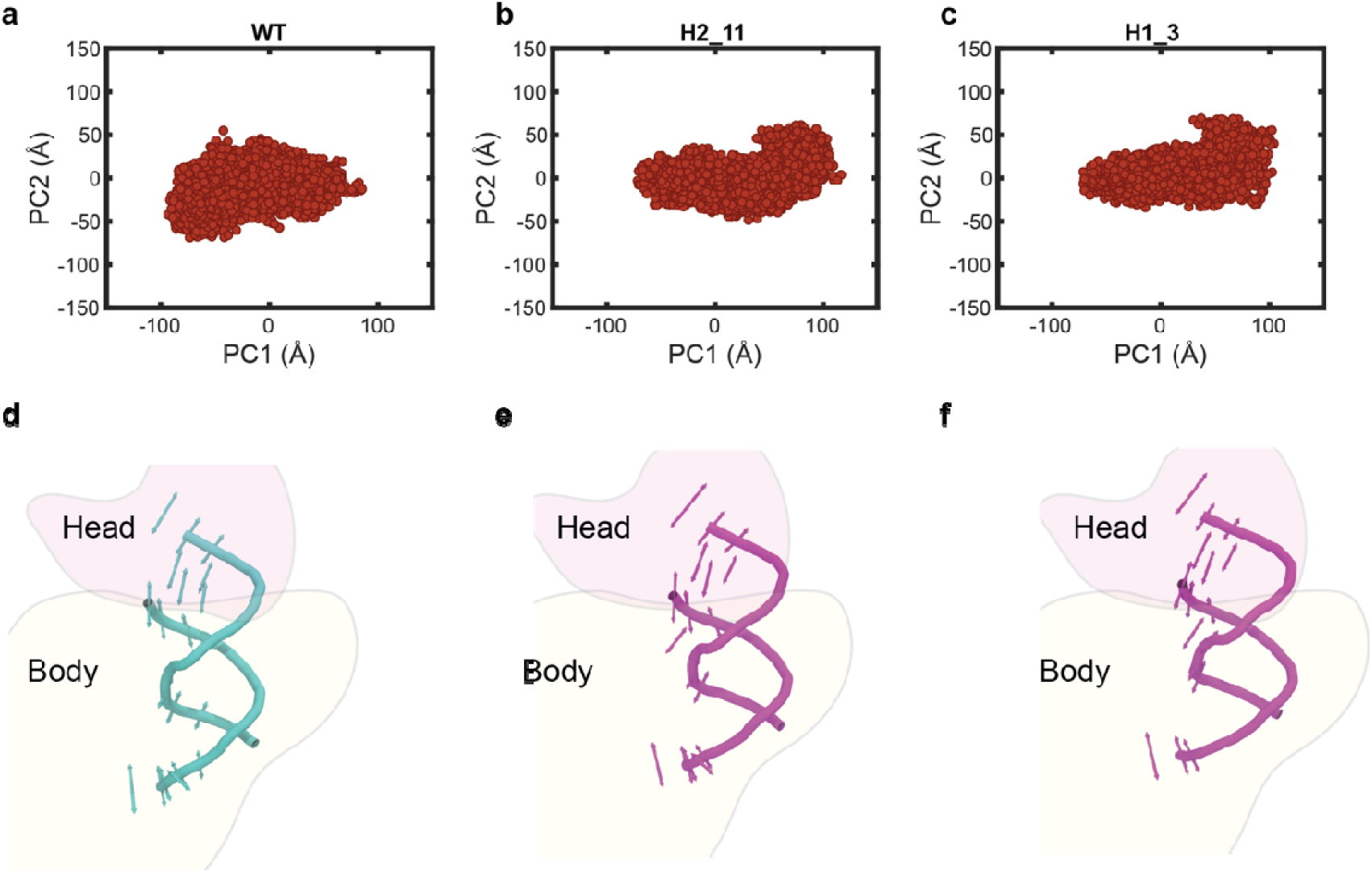
Hinge 1 variants produce a more compact hinge 1 during forward swivel motions. 2 dimensional projections of principal component 1 (PC1) and 2 (PC2) for (**a**) WT, and the (**b**) H2_11, (**c**) H1_3 hinge variants. Structural description of PC1 projected onto the structure of hinge 1 of the SSU for (**d**) WT, and the (**e**) H2_11, (**f**) H1_3 hinge variants.

**Figure S3.**
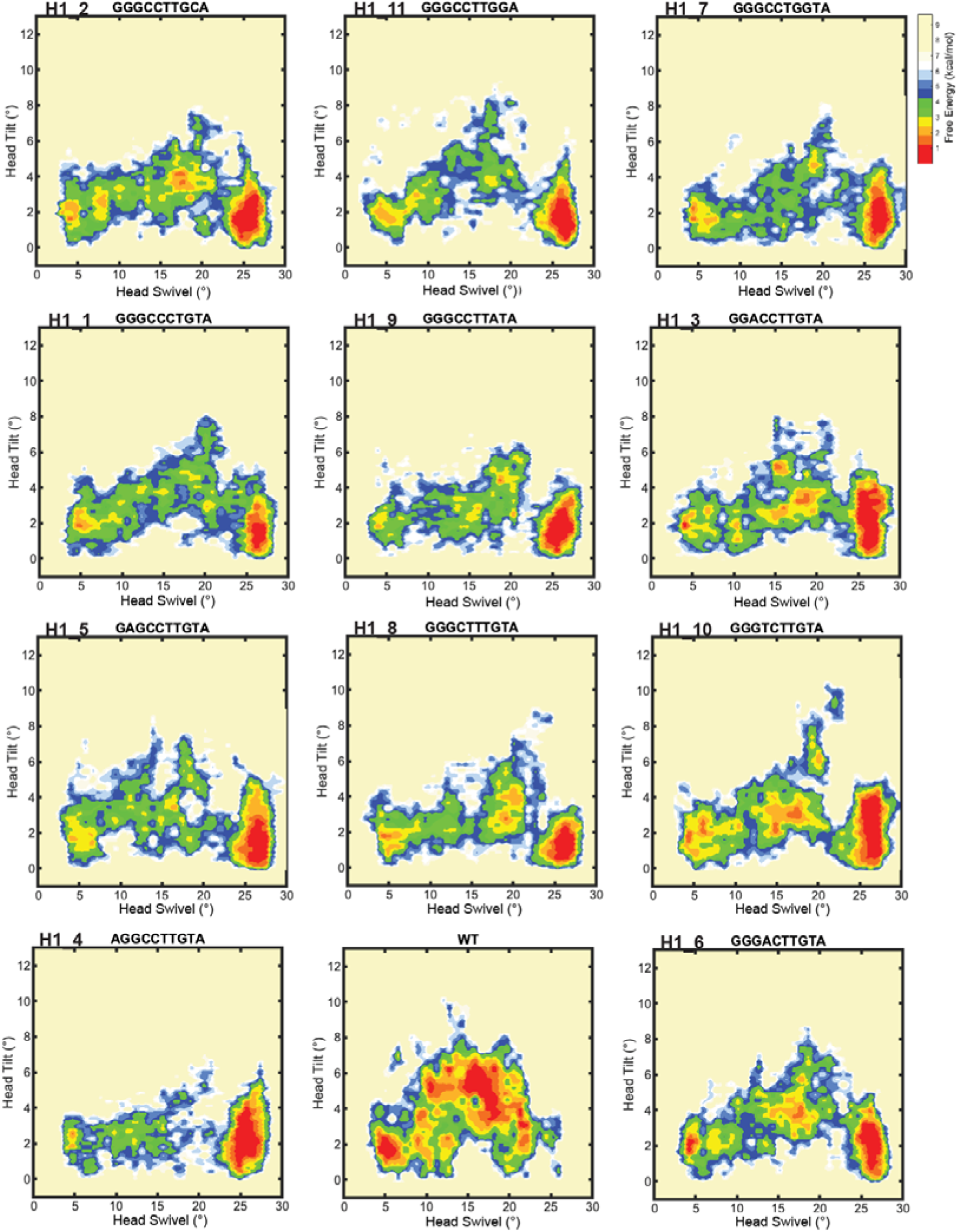
Head domain motion in forward swivel for hinge 1 variants. Free-energy landscapes of the forward swivel (PRE to INT3) stage of translocation with head domain swivel and tilt angles as reaction coordinates. The landscapes are based on Boltzmann weighted probability distributions. The heat map is a normalized number of frames in the targeted MD simulation, where the frame count is divided by the maximum frame count. Shown are the maps for the top 12 hinge 1 enriched genotypes. Maps correspond to ten trajectories per genotype.

**Figure S4.**
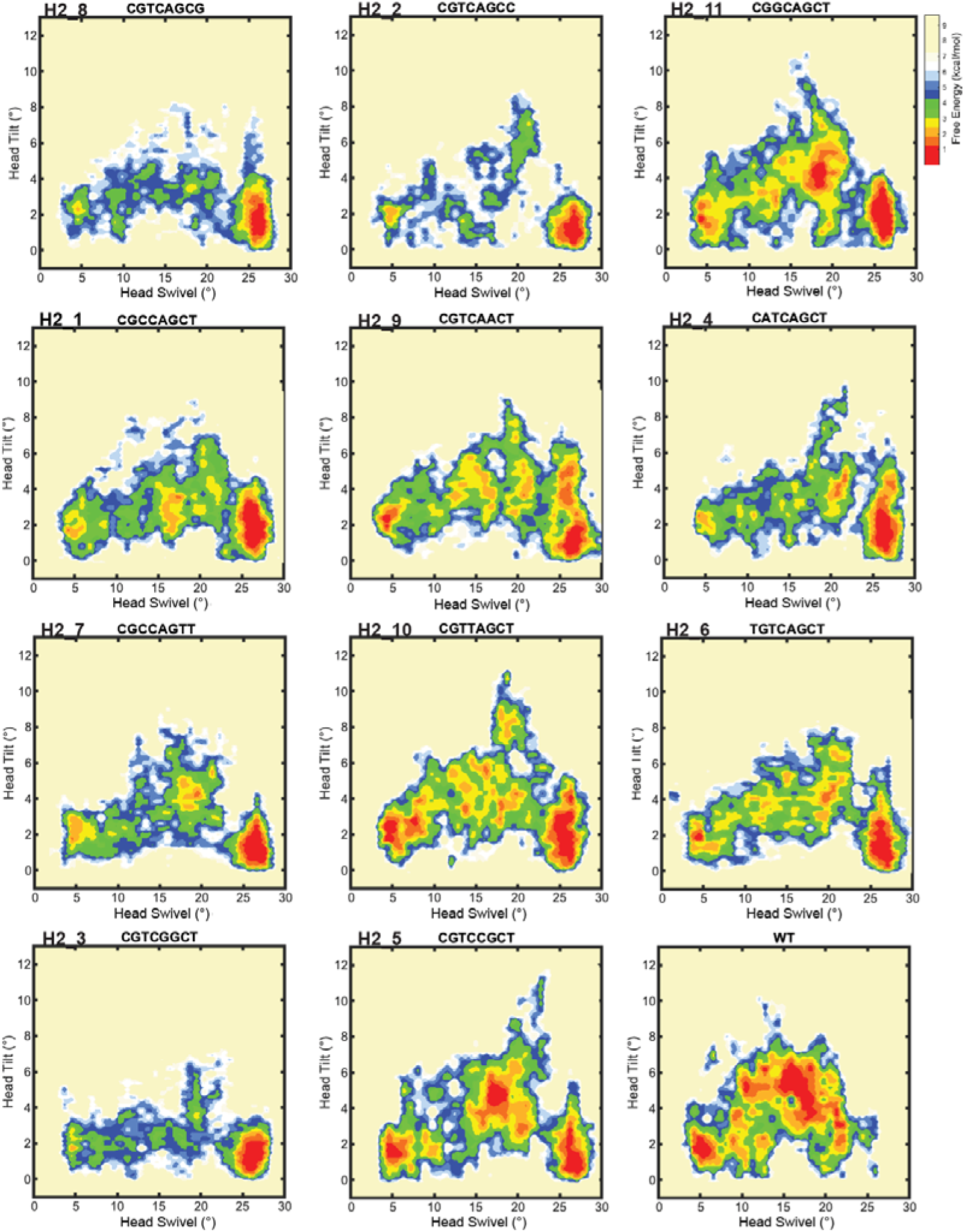
Head domain motion in forward swivel for hinge 2 variants. Free-energy landscapes of the forward swivel (PRE to INT3) stage of translocation with head domain swivel and tilt angles as reaction coordinates. The landscapes are based on Boltzmann weighted probability distributions. The heat map is a normalized number of frames in the targeted MD simulation, where the frame count is divided by the maximum frame count. Shown are the maps for the top 12 hinge 2 enriched genotypes. Maps correspond to ten trajectories per genotype.

**Figure S5.**
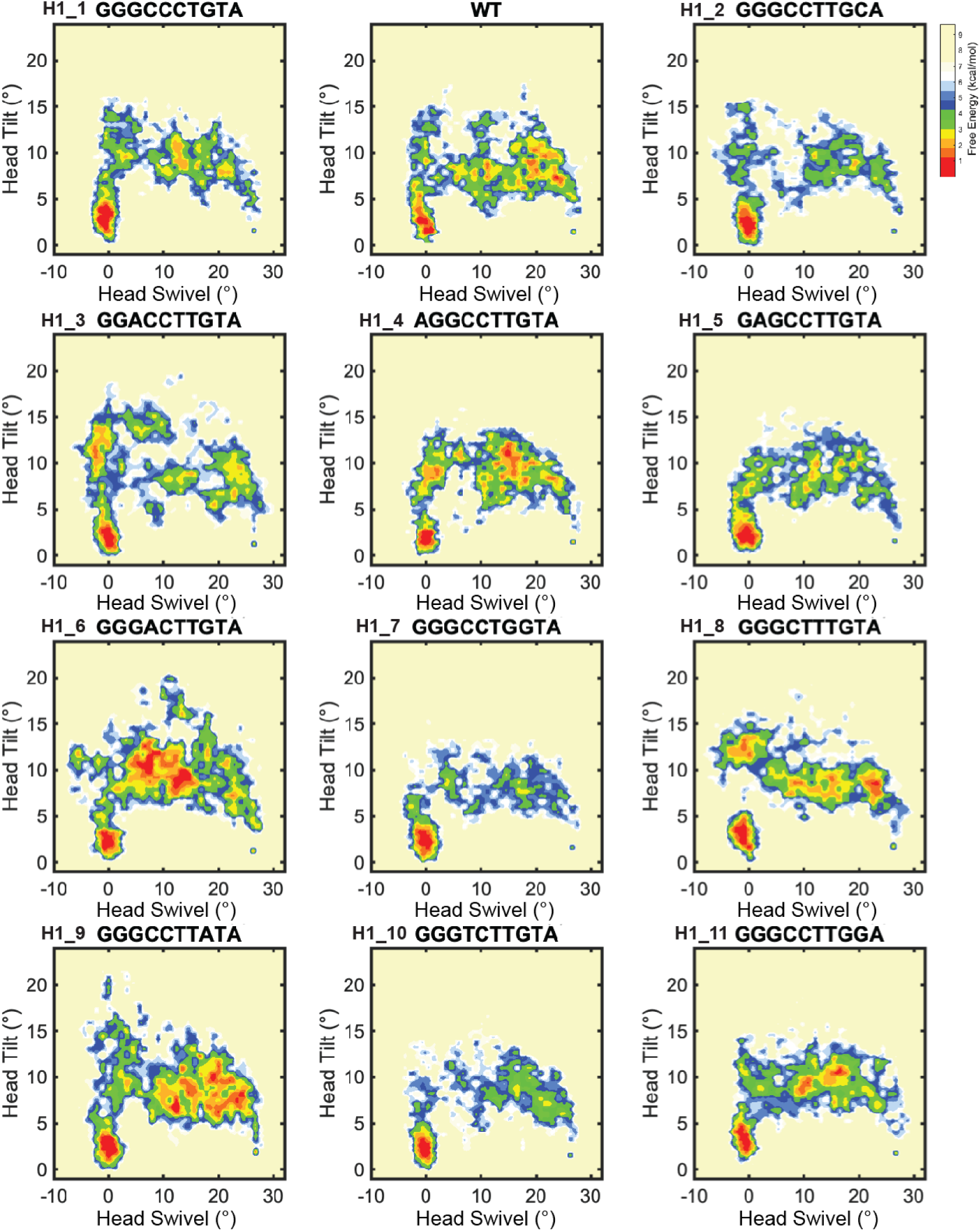
Head domain motion in reverse swivel for hinge 1 variants. Free-energy landscapes of the reverse swivel (INT3 to POST) stage of translocation with head domain swivel and tilt angles as reaction coordinates. The landscapes are based on Boltzmann weighted probability distributions. The heat map is a normalized number of frames in the targeted MD simulation, where the frame count is divided by the maximum frame count. Shown are the maps for the top 12 hinge 1 enriched genotypes. Maps correspond to ten trajectories per genotype.

**Figure S6.**
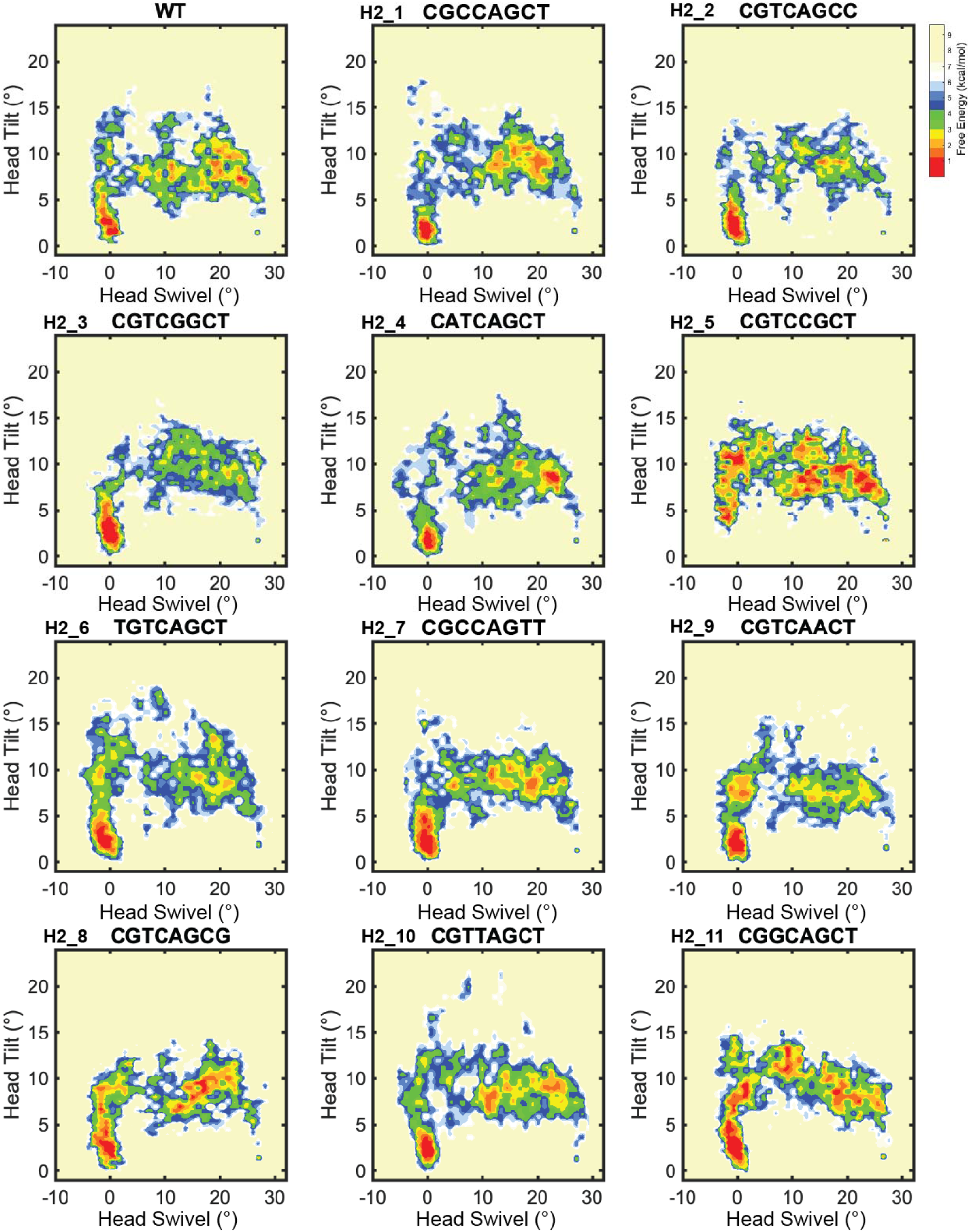
Head domain motion in reverse swivel for hinge 2 variants. Free-energy landscapes of the reverse swivel (INT3 to POST) stage of translocation with head domain swivel and tilt angles as reaction coordinates. The landscapes are based on Boltzmann weighted probability distributions. The heat map is a normalized number of frames in the targeted MD simulation, where the frame count is divided by the maximum frame count. Shown are the maps for the top 12 hinge 2 enriched genotypes. Maps correspond to ten trajectories per genotype.

**Figure S7.**
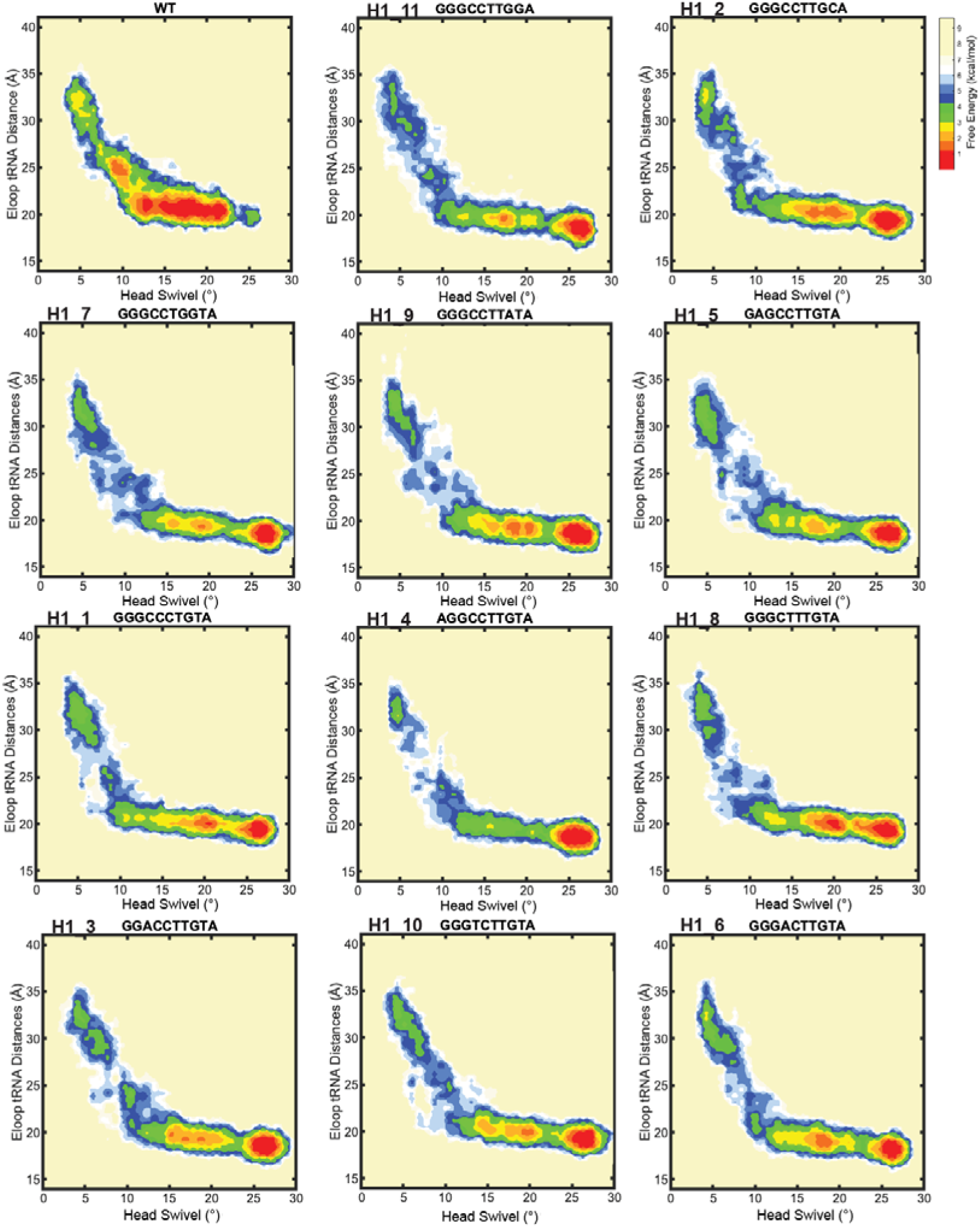
Deacyl tRNA motion in forward swivel for hinge 1 variants. Free-energy landscapes of the forward swivel (PRE to INT3) stage of translocation with head domain swivel angle and E loop-Deacyl tRNA distance as reaction coordinates. The landscapes are based on Boltzmann weighted probability distributions. The heat map is a normalized number of frames in the targeted MD simulation, where the frame count is divided by the maximum frame count. Shown are the maps for the top 12 hinge 1 enriched genotypes. Maps correspond to ten trajectories per genotype.

**Figure S8.**
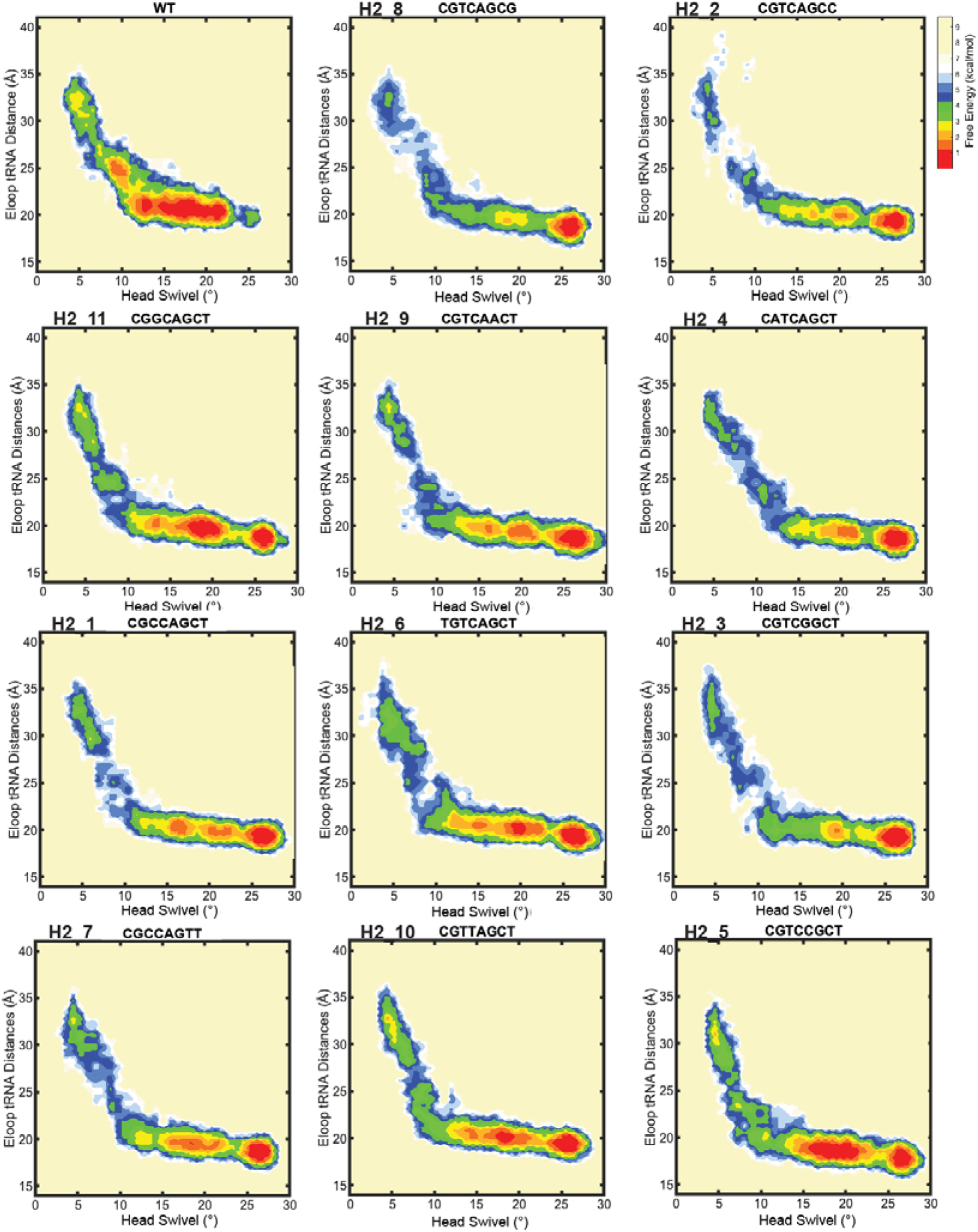
Deacyl tRNA motion in forward swivel for hinge 2 variants. Free-energy landscapes of the forward swivel (PRE to INT3) stage of translocation with head domain swivel angle and E loop-Deacyl tRNA distance as reaction coordinates. The landscapes are based on Boltzmann weighted probability distributions. The heat map is a normalized number of frames in the targeted MD simulation, where the frame count is divided by the maximum frame count. Shown are the maps for the top 12 hinge 2 enriched genotypes. Maps correspond to ten trajectories per genotype.

**Figure S9.**
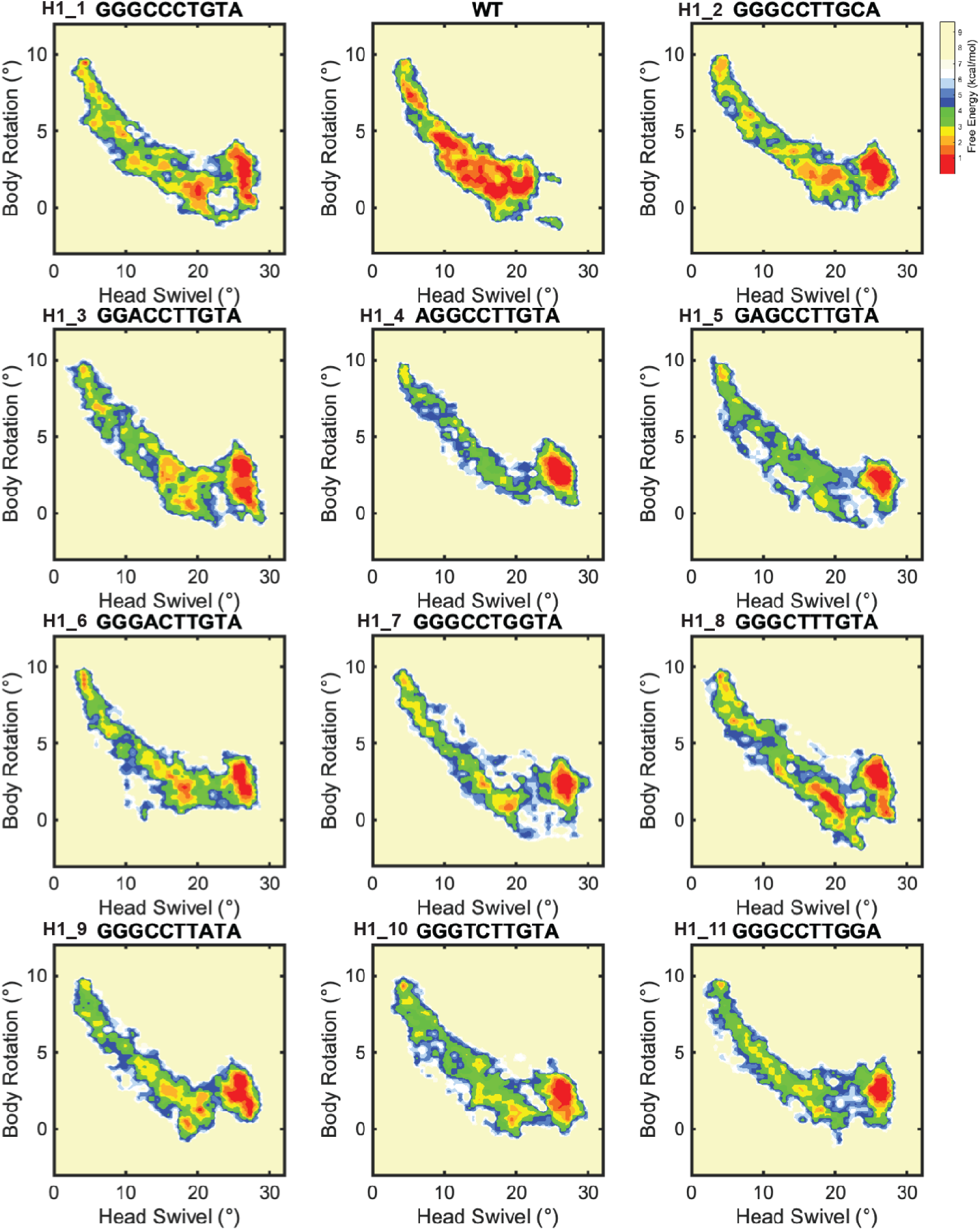
Small subunit rotation in forward swivel for hinge 1 variants. Free-energy landscapes of the forward swivel (PRE to INT3) stage of translocation with head domain swivel angle and small subunit rotation as reaction coordinates. The landscapes are based on Boltzmann weighted probability distributions. The heat map is a normalized number of frames in the targeted MD simulation, where the frame count is divided by the maximum frame count. Shown are the maps for the top 12 hinge 1 enriched genotypes. Maps correspond to ten trajectories per genotype.

**Figure S10.**
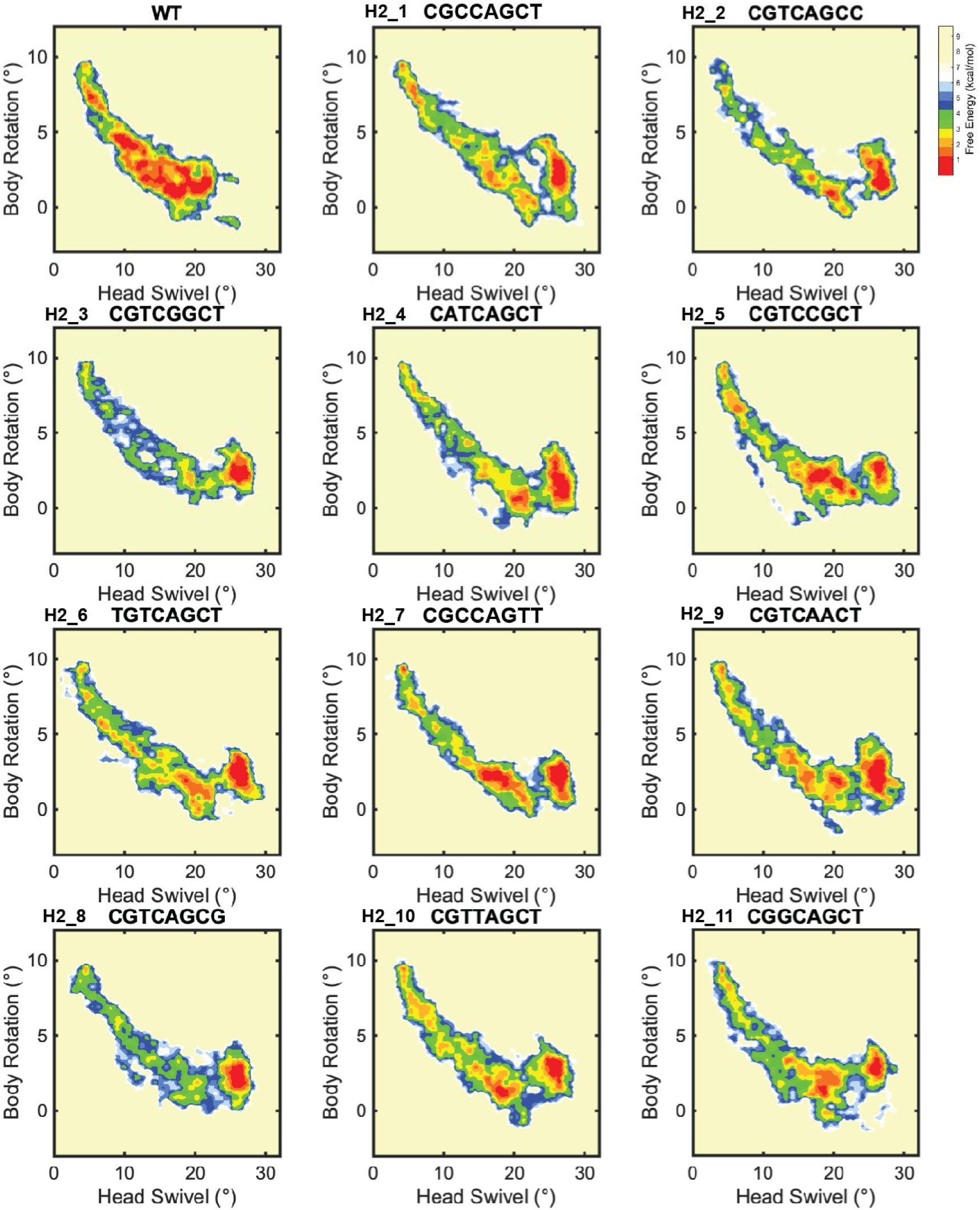
Small subunit rotation in forward swivel for hinge 2 variants. Free-energy landscapes of the forward swivel (PRE to INT3) stage of translocation with head domain swivel angle and small subunit rotation as reaction coordinates. The landscapes are based on Boltzmann weighted probability distributions. The heat map is a normalized number of frames in the targeted MD simulation, where the frame count is divided by the maximum frame count. Shown are the maps for the top 12 hinge 2 enriched genotypes. Maps correspond to ten trajectories per genotype.

**Table S1.**
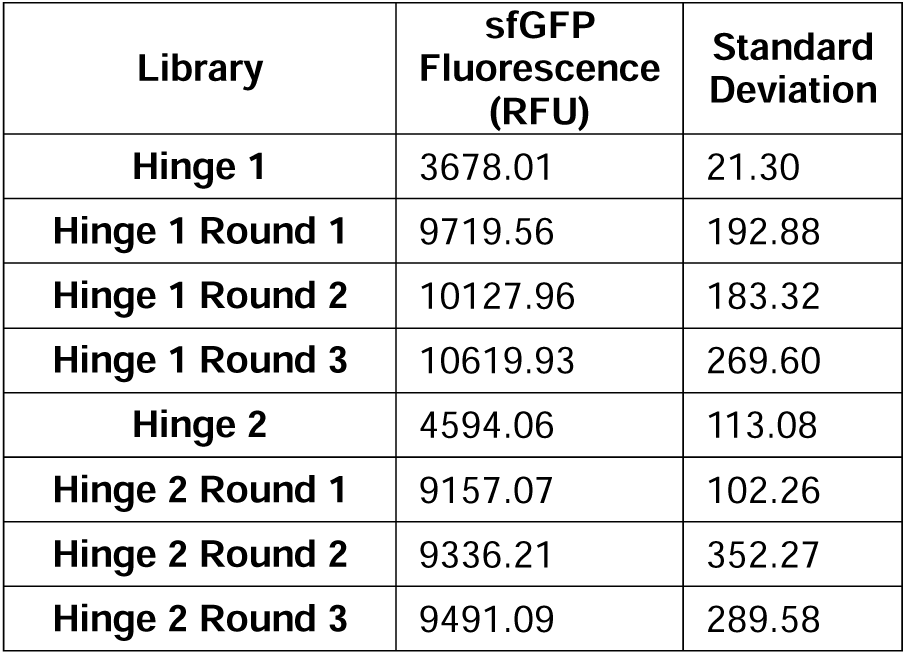
Activity of hinge libraries. The activity of hinge 1 and hinge 2 libraries across rounds of RISE (Ribosome synthesis and evolution) was quantified by sfGFP production in iSAT (*in vitro* synthesis and translation). Standard deviations were obtained from three independent experiments.

| Library | sfGFP Fluorescence (RFU) | Standard Deviation |
| --- | --- | --- |
| Hinge 1 | 3678.01 | 21.30 |
| Hinge 1 Round 1 | 9719.56 | 192.88 |
| Hinge 1 Round 2 | 10127.96 | 183.32 |
| Hinge 1 Round 3 | 10619.93 | 269.60 |
| Hinge 2 | 4594.06 | 113.08 |
| Hinge 2 Round 1 | 9157.07 | 102.26 |
| Hinge 2 Round 2 | 9336.21 | 352.27 |
| Hinge 2 Round 3 | 9491.09 | 289.58 |

**Table S2.** Top RISE-enriched hinge variants. Top genotypes by enrichment (reads in final pool relative to reads in initial library) for hinge libraries after three rounds of RISE. Mutations relative to wild type are shown in red.

| Hinge 1 Library Selection |  |  | Hinge 2 Library Selection |  |  |
| --- | --- | --- | --- | --- | --- |
| Name | Genotype | Enrichment | Name | Genotype | Enrichment |
| H1_1 | GGGCCCTGTA | 1554.837919 | WT | CGTCAGCT | 2210.154204 |
| WT | GGGCCTTGTA | 617.2618836 | H2_1 | CGCCAGCT | 1636.165946 |
| H1_2 | GGGCCTTGCA | 309.747947 | H2_2 | CGTCAGCC | 420.5490476 |
| H1_3 | GGACCTTGTA | 267.6572245 | H2_3 | CGTCGGCT | 96.18733749 |
| H1_4 | AGGCCTTGTA | 215.3586732 | H2_4 | CATCAGCT | 27.74690234 |
| H1_5 | GAGCCTTGTA | 182.547795 | H2_5 | CGTCCGCT | 14.04559224 |
| H1_6 | GGGACTTGTA | 46.9294985 | H2_6 | TGTCAGCT | 13.71668665 |
| H1_7 | GGGCCTGGTA | 42.28957633 | H2_7 | CGTCAGTT | 11.8496328 |
| H1_8 | GGGCTTTGTA | 38.97534621 | H2_8 | CGTCAGCG | 10.18725777 |
| H1_9 | GGGCCTTATA | 38.25947251 | H2_9 | CGTCAACT | 10.06832335 |
| H1_10 | GGGTCTTGTA | 19.1893924 | H2_10 | CGTTAGCT | 9.837163922 |
| H1_11 | GGGCCTTGGA | 18.13546722 | H2_11 | CGGCAGCT | 4.343979738 |

**Table S3.** Simulation transition times for forward and reverse head swivel. The average times across simulated trajectories for the transitions PRE to 10° head domain swivel, PRE to INT2, PRE to INT3, and INT3 to POST, calculated as number of frames to achieve each destination state. *t_ru_*is one reduced time unit.

| Variant | PRE to 10° ($t_{ru}$) | SD | PRE to INT2 ($t_{ru}$) | SD | PRE to INT3 ($t_{ru}$) | SD | INT3 to POST ($t_{ru}$) | SD |
| --- | --- | --- | --- | --- | --- | --- | --- | --- |
| WT | 140.3 | 51.3 | 679.4 | 215.7 | 915.6 | 161.0 | 272.0 | 87.9 |
| H1_1 | 158.2 | 66.3 | 531.8 | 208.4 | 571.1 | 201.4 | 812.7 | 433.0 |
| H1_2 | 159.5 | 44.4 | 491.2 | 78.3 | 531.8 | 300.1 | 498 | 357.2 |
| H1_3 | 118.5 | 38.3 | 417.6 | 84.3 | 494.7 | 159.7 | 411.7 | 476.1 |
| H1_4 | 99.4 | 22.5 | 394.4 | 41.2 | 391.6 | 69.5 | 532.8 | 360.7 |
| H1_5 | 145.4 | 58.1 | 500.6 | 149.9 | 497.1 | 190.6 | 392.1 | 216.9 |
| H1_6 | 140 | 41.0 | 497.0 | 113.0 | 536.0 | 147.5 | 578.2 | 520.9 |
| H1_7 | 141.2 | 43.6 | 447.2 | 94.9 | 471.2 | 100.8 | 345.1 | 235.6 |
| H1_8 | 144.3 | 35.6 | 472.6 | 96.8 | 567.3 | 163.6 | 423.3 | 313.0 |
| H1_9 | 120.6 | 30.5 | 436.8 | 82.7 | 496.7 | 159.1 | 343.4 | 284.3 |
| H1_10 | 136.5 | 58.3 | 525.4 | 127.5 | 503.2 | 147.7 | 515.9 | 460.4 |
| H2_1 | 130.0 | 20.9 | 423.6 | 56.3 | 521.8 | 106.5 | 567.8 | 308.6 |
| H2_2 | 119.3 | 29.2 | 442.0 | 44.0 | 534.4 | 300.1 | 498 | 357.2 |
| H2_3 | 125.9 | 33.3 | 440.4 | 79.7 | 464.5 | 73.2 | 417.4 | 158.6 |
| H2_4 | 97.5 | 23.1 | 375.6 | 68.9 | 476.4 | 101.0 | 564.9 | 360.1 |
| H2_5 | 151.3 | 48.6 | 464.4 | 93.5 | 625.1 | 159.2 | 690.5 | 230.3 |
| H2_6 | 143.3 | 26.2 | 505.6 | 86.6 | 584.3 | 210.6 | 506.1 | 332.3 |
| H2_7 | 128.2 | 22.9 | 461.4 | 99.1 | 544.2 | 157.6 | 493.6 | 378.8 |
| H2_8 | 112.2 | 40.8 | 421.6 | 135.0 | 459.3 | 100.7 | 582.4 | 358.8 |
| H2_9 | 127.8 | 44.7 | 426.0 | 84.2 | 526.2 | 156.0 | 526.3 | 408.6 |
| H2_10 | 156.5 | 42.3 | 525.4 | 116.9 | 615.6 | 188.7 | 545.3 | 419.9 |

## References

1. Katunin, V.I., A. Savelsbergh, M. V. Rodnina, and W. Wintermeyer. 2002. Coupling of GTP hydrolysis by elongation factor G to translocation and factor recycling on the ribosome. Biochemistry. 41:12806–12812.

2. Fredrick, K. 2003. Catalysis of Ribosomal Translocation by Sparsomycin. Science. 300:1159–1162.

3. Gavrilova, L.P., O.E. Kostiashkina, V.E. Koteliansky, N.M. Rutkevitch, and A.S. Spirin. 1976. Factor-free (“Non-enzymic”) and factor-dependent systems of translation of polyuridylic acid by Escherichia coli ribosomes. J. Mol. Biol. 101:537–552.

4. Shoji, S., S.E. Walker, and K. Frederick. 2009. Ribosomal translocation: one step closer to the molecular mechanism. ACS Chem. Biol. 4:93–107.

5. Agirrezabala, X., and J. Frank. 2009. Elongation in translation as a dynamic interaction among the ribosome, tRNA, and elongation factors EF-G and EF-Tu. Q. Rev. Biophys. 42:159–200.

6. Ratje, A.H., J. Loerke, A. Mikolajka, M. Brünner, P.W. Hildebrand, A.L. Starosta, A. Dönhöfer, S.R. Connell, P. Fucini, T. Mielke, P.C. Whitford, J.N. Onuchic, Y. Yu, K.Y. Sanbonmatsu, R.K. Hartmann, P.A. Penczek, D.N. Wilson, and C.M.T. Spahn. 2010. Head swivel on the ribosome facilitates translocation by means of intra-subunit tRNA hybrid sites. Nature. 468:713–716.

7. Rundlet, E.J., M. Holm, M. Schacherl, S.K. Natchiar, R.B. Altman, C.M.T. Spahn, A.G. Myasnikov, and S.C. Blanchard. 2021. Structural basis of early translocation events on the ribosome. Nature. 595:741–745.

8. Zhou, J., L. Lancaster, J.P. Donohue, and H.F. Noller. 2014. How the ribosome hands the A-site tRNA to the P site during EF-G-catalyzed translocation. Science. 345:1188–1191.

9. Ramrath, D.J.F., L. Lancaster, T. Sprink, T. Mielke, J. Loerke, H.F. Noller, and C.M.T. Spahn. 2013. Visualization of two transfer RNAs trapped in transit during elongation factor G-mediated translocation. Proc. Natl. Acad. Sci. U. S. A. 110:20964–20969.

10. Brilot, A.F., A.A. Korostelev, D.N. Ermolenko, and N. Grigorieff. 2013. Structure of the ribosome with elongation factor G trapped in the pretranslocation state. Proc. Natl. Acad. Sci. U. S. A. 110:20994–20999.

11. Gao, Y.-G., M. Selmer, C.M. Dunham, A. Weixlbaumer, A.C. Kelley, and V. Ramakrishnan. 2009. The Structure of the Ribosome with Elongation Factor G Trapped in the Posttranslocational State. Science. 326:694–699.

12. Guo, Z., and H.F. Noller. 2012. Rotation of the head of the 30S ribosomal subunit during mRNA translocation. Proc. Natl. Acad. Sci. U. S. A. 109:20391–20394.

13. Wilden, B., A. Savelsbergh, M. V Rodnina, and W. Wintermeyer. 2006. Role and timing of GTP binding and hydrolysis during EF-G-dependent tRNA translocation on the ribosome. Proc. Natl. Acad. Sci. U. S. A. 103:13670–13675.

14. Ermolenko, D.N., and H.F. Noller. 2011. mRNA translocation occurs during the second step of ribosomal intersubunit rotation. Nat. Struct. Mol. Biol. 18:457–463.

15. Wasserman, M.R., J.L. Alejo, R.B. Altman, and S.C. Blanchard. 2016. Multiperspective smFRET reveals rate-determining late intermediates of ribosomal translocation. Nat. Struct. Mol. Biol. 23:333–341.

16. Alejo, J.L., and S.C. Blanchard. 2017. Miscoding-induced stalling of substrate translocation on the bacterial ribosome. Proc. Natl. Acad. Sci. U. S. A. 114:E8603–E8610.

17. Munro, J.B., M.R. Wasserman, R.B. Altman, L. Wang, and S.C. Blanchard. 2010. Correlated conformational events in EF-G and the ribosome regulate translocation. Nat. Struct. Mol. Biol. 17:1470–1477.

18. Savelsbergh, A., V.I. Katunin, D. Mohr, F. Peske, M. V Rodnina, and W. Wintermeyer. 2003. An Elongation Factor G-Induced Ribosome Rearrangement Precedes tRNA-mRNA Translocation. Mol. Cell. 11:1517–1523.

19. Borovinskaya, M.A., S. Shoji, J.M. Holton, K. Fredrick, and J.H.D. Cate. 2007. A steric block in translation caused by the antibiotic spectinomycin. ACS Chem. Biol. 2:545–552.

20. Nishima, W., D. Girodat, M. Holm, E.J. Rundlet, J.L. Alejo, K. Fischer, S.C. Blanchard, and K.Y. Sanbonmatsu. 2022. Hyper-swivel head domain motions are required for complete mRNA-TRNA translocation and ribosome resetting. Nucleic Acids Res. 50:8302–8320.

21. Milicevic, N., L. Jenner, A. Myasnikov, M. Yusupov, and G. Yusupova. 2023. mRNA reading frame maintenance during eukaryotic ribosome translocation. Nature.

22. Bock, L. V, C. Blau, G.F. Schröder, I.I. Davydov, N. Fischer, H. Stark, M. V Rodnina, A.C. Vaiana, and H. Grubmüller. 2013. Energy barriers and driving forces in tRNA translocation through the ribosome. Nat. Struct. Mol. Biol. 20:1390–1396.

23. Nguyen, K., and P.C. Whitford. 2016. Steric interactions lead to collective tilting motion in the ribosome during mRNA-tRNA translocation. Nat. Commun. 7:1–9.

24. Flis, J., M. Holm, E.J. Rundlet, J. Loerke, T. Hilal, M. Dabrowski, J. Bürger, T. Mielke, S.C. Blanchard, C.M.T. Spahn, and T. V Budkevich. 2018. tRNA Translocation by the Eukaryotic 80S Ribosome and the Impact of GTP Hydrolysis. Cell Rep. 25:2676–2688.e7.

25. Hassan, A., S. Byju, F.C. Freitas, C. Roc, N. Pender, K. Nguyen, E.M. Kimbrough, J.M. Mattingly, R.L. Gonzalez, R.J. De Oliveira, C.M. Dunham, and P.C. Whitford. 2023. Ratchet, swivel, tilt and roll: a complete description of subunit rotation in the ribosome. Nucleic Acids Res. 51:919–934.

26. Mohan, S., J.P. Donohue, and H.F. Noller. 2014. Molecular mechanics of 30S subunit head rotation. Proc. Natl. Acad. Sci. 111:13325–13330.

27. Noller, H.F., J.P. Donohue, and R.R. Gutell. 2022. The universally conserved nucleotides of the small subunit ribosomal RNAs. RNA. 28:623–644.

28. Hammerling, M.J., B.R. Fritz, D.J. Yoesep, D.S. Kim, E.D. Carlson, and M.C. Jewett. 2020. In vitro ribosome synthesis and evolution through ribosome display. Nat. Commun. 11:1–10.

29. Whitford, P.C., S.C. Blanchard, J.H.D. Cate, and K.Y. Sanbonmatsu. 2013. Connecting the Kinetics and Energy Landscape of tRNA Translocation on the Ribosome. PLoS Comput. Biol. 9.

30. Jewett, M.C., B.R. Fritz, L.E. Timmerman, and G.M. Church. 2013. In vitro integration of ribosomal RNA synthesis, ribosome assembly, and translation. Mol. Syst. Biol. 9:1–8.

31. D’Aquino, A.E., T. Azim, N.A. Aleksashin, A.J. Hockenberry, A. Krüger, and M.C. Jewett. 2020. Mutational characterization and mapping of the 70S ribosome active site. Nucleic Acids Res. 48:2777–2789.

32. Kofman, C., A.M. Watkins, D.S. Kim, J.A. Willi, A.C. Wooldredge, A.S. Karim, R. Das, and M.C. Jewett. 2022. Computationally-guided design and selection of high performing ribosomal active site mutants. Nucleic Acids Res. 50:13143–13154.

33. Krüger, A., A.M. Watkins, R. Wellington-Oguri, J. Romano, C. Kofman, A. DeFoe, Y. Kim, J. Anderson-Lee, E. Fisker, J. Townley, A.E. D’Aquino, R. Das, and M.C. Jewett. 2023. Community science designed ribosomes with beneficial phenotypes. Nat. Commun. 14:961.

34. Asai, T., D. Zaporojets, C. Squires, and C.L. Squires. 1999. An escherichia coli strain with all chromosomal rRNA operons inactivated: Complete exchange of rRNA genes between bacteria. Proc. Natl. Acad. Sci. U. S. A. 96:1971–1976.

35. Carlson, E.D., A.E. d’Aquino, D.S. Kim, E.M. Fulk, K. Hoang, T. Szal, A.S. Mankin, and M.C. Jewett. 2019. Engineered ribosomes with tethered subunits for expanding biological function. Nat. Commun. 10:1–13.

36. Dunkle, J.A., L. Wang, M.B. Feldman, A. Pulk, V.B. Chen, G.J. Kapral, J. Noeske, J.S. Richardson, S.C. Blanchard, and J.H.D. Cate. 2011. Structures of the bacterial ribosome in classical and hybrid states of tRNA binding. Science. 332:981–984.

37. Zhou, J., L. Lancaster, J.P. Donohue, and H.F. Noller. 2013. Crystal structures of EF-G - Ribosome complexes trapped in intermediate states of translocation. Science. 340:1236086.

38. Guyomar, C., G. D’Urso, S. Chat, E. Giudice, and R. Gillet. 2021. Structures of tmRNA and SmpB as they transit through the ribosome. Nat. Commun. 12:4909.

39. Hoffer, E.D., S. Hong, S. Sunita, T. Maehigashi, R.L. Gonzalez Jnr, P.C. Whitford, and C.M. Dunham. 2020. Structural insights into mRNA reading frame regulation by tRNA modification and slippery codon–anticodon pairing. Elife. 9:1–20.

40. Shah, P., D.M. McCandlish, and J.B. Plotkin. 2015. Contingency and entrenchment in protein evolution under purifying selection. Proc. Natl. Acad. Sci. U. S. A. 112:E3226–E3235.

41. Petrov, A.S., B. Gulen, A.M. Norris, N.A. Kovacs, C.R. Bernier, K.A. Lanier, G.E. Fox, S.C. Harvey, R.M. Wartell, N. V. Hud, and L.D. Williams. 2015. History of the ribosome and the origin of translation. Proc. Natl. Acad. Sci. U. S. A. 112:15396– 15401.

42. Basan, M., T. Honda, D. Christodoulou, M. Hörl, Y.F. Chang, E. Leoncini, A. Mukherjee, H. Okano, B.R. Taylor, J.M. Silverman, C. Sanchez, J.R. Williamson, J. Paulsson, T. Hwa, and U. Sauer. 2020. A universal trade-off between growth and lag in fluctuating environments. Nature. 584:470–474.

43. Espinosa, R., M.A. Sørensen, and S. Lo Svenningsen. 2022. Escherichia coli protein synthesis is limited by mRNA availability rather than ribosomal capacity during phosphate starvation. Front. Microbiol. 13:989818.

44. Farewell, A., and F.C. Neidhardt. 1998. Effect of temperature on in vivo protein synthetic capacity in Escherichia coli. J. Bacteriol. 180:4704–4710.

45. Fritz, B.R., and M.C. Jewett. 2014. The impact of transcriptional tuning on in vitro integrated rRNA transcription and ribosome construction. Nucleic Acids Res. 42:6774–6785.

46. Fritz, B.R., O.K. Jamil, and M.C. Jewett. 2015. Implications of macromolecular crowding and reducing conditions for in vitro ribosome construction. Nucleic Acids Res. 43:4774–4784.

47. Huang, S., N.A. Aleksashin, A.B. Loveland, D. Klepacki, K. Reier, A. Kefi, T. Szal, J. Remme, L. Jaeger, N. Vázquez-Laslop, A.A. Korostelev, and A.S. Mankin. 2020. Ribosome engineering reveals the importance of 5S rRNA autonomy for ribosome assembly. Nat. Commun. 11:1–13.

48. Phillips, J.C., R. Braun, W. Wang, J. Gumbart, E. Tajkhorshid, E. Villa, C. Chipot, R.D. Skeel, L. Kalé, and K. Schulten. 2005. Scalable molecular dynamics with NAMD. J. Comput. Chem. 26:1781–1802.

49. Best, R.B., X. Zhu, J. Shim, P.E.M. Lopes, J. Mittal, M. Feig, and A.D. MacKerell. 2012. Optimization of the additive CHARMM all-atom protein force field targeting improved sampling of the backbone φ, ψ and side-chain χ1 and χ2 Dihedral Angles. J. Chem. Theory Comput. 8:3257–3273.

50. Pettersen, E.F., T.D. Goddard, C.C. Huang, G.S. Couch, D.M. Greenblatt, E.C. Meng, and T.E. Ferrin. 2004. UCSF Chimera - A visualization system for exploratory research and analysis. J. Comput. Chem. 25:1605–1612.

51. Pronk, S., S. Páll, R. Schulz, P. Larsson, P. Bjelkmar, R. Apostolov, M.R. Shirts, J.C. Smith, P.M. Kasson, D. Van Der Spoel, B. Hess, and E. Lindahl. 2013. GROMACS 4.5: A high-throughput and highly parallel open source molecular simulation toolkit. Bioinformatics. 29:845–854.

52. Abraham, M.J., T. Murtola, R. Schulz, S. Páll, J.C. Smith, B. Hess, and E. Lindah. 2015. Gromacs: High performance molecular simulations through multi-level parallelism from laptops to supercomputers. SoftwareX. 1–2:19–25.

53. Whitford, P.C., J.K. Noel, S. Gosavi, A. Schug, K.Y. Sanbonmatsu, and J.N. Onuchic. 2009. An all-atom structure-based potential for proteins: Bridging minimal models with all-atom empirical forcefields. Proteins Struct. Funct. Bioinforma. 75:430–441.

54. Girodat, D.J., H.-J. Wieden, S.C. Blanchard, and K.Y. Sanbonmatsu. 2023. Geometric alignment of aminoacyl-tRNA relative to catalytic centers of the ribosome underpins accurate mRNA decoding. Biophys. J. 122:488a.

55. Yang, Y.I., Q. Shao, J. Zhang, L. Yang, and Y.Q. Gao. 2019. Enhanced sampling in molecular dynamics. J. Chem. Phys. 151:070902.

56. Lavery, R., J.H. Maddocks, M. Pasi, and K. Zakrzewska. 2014. Analyzing ion distributions around DNA. Nucleic Acids Res. 42:8138–8149.

57. Blanchet, C., M. Pasi, K. Zakrzewska, and R. Lavery. 2011. CURVES+ web server for analyzing and visualizing the helical, backbone and groove parameters of nucleic acid structures. Nucleic Acids Res. 39:68–73.

